# Interpretable models for scRNA-seq data embedding with multi-scale structure preservation

**DOI:** 10.1101/2023.11.23.568428

**Authors:** David Novak, Cyril de Bodt, Pierre Lambert, John Aldo Lee, Sofie Van Gassen, Yvan Saeys

## Abstract

The ability to explore high-dimensional single-cell transcriptomics data efficiently is crucial in many biological studies. Dimensionality reduction techniques have therefore emerged as a basic building block of analytical workflows. They generate low-dimensional embeddings that capture important structures in the data, and are often used in discovery, quality control, and downstream analysis. However, the trustworthiness of current methods and the rigour of popular evaluation criteria are limited. We tackle this in an empirical study of structure-preserving data embeddings, delivering two tools. First, we introduce *ViScore*: a robust scoring framework that improves both unsupervised and supervised quality metrics, with emphasis on scalability and fairness. Second, we introduce *ViVAE* : a deep learning model that achieves better multi-scale structure preservation and is equipped with new tools for interpretability. We demonstrate the potential of these contributions to advance the trustworthiness of single-cell dimensionality reduction in a quantitative comparison and focused case studies.

## INTRODUCTION

Data embedding using dimensionality reduction (DR) is common in the computational analysis of single-cell RNA-sequencing (scRNA-seq) data^1,2^. DR converts high-dimensional (HD) gene expression data to low-dimensional (LD) embeddings (Figure 1**A**). These embeddings are suitable for describing cell type diversity^2^, quality control (QC) after data integration^3^, or representing tentative trajectory structures^4^. They generally improve the interpretability of single-cell datasets, which are sparse and noise-prone^5^. DR is therefore crucial for effective exploratory data analysis in the single-cell bioinformatics domain.

**Figure 1.**
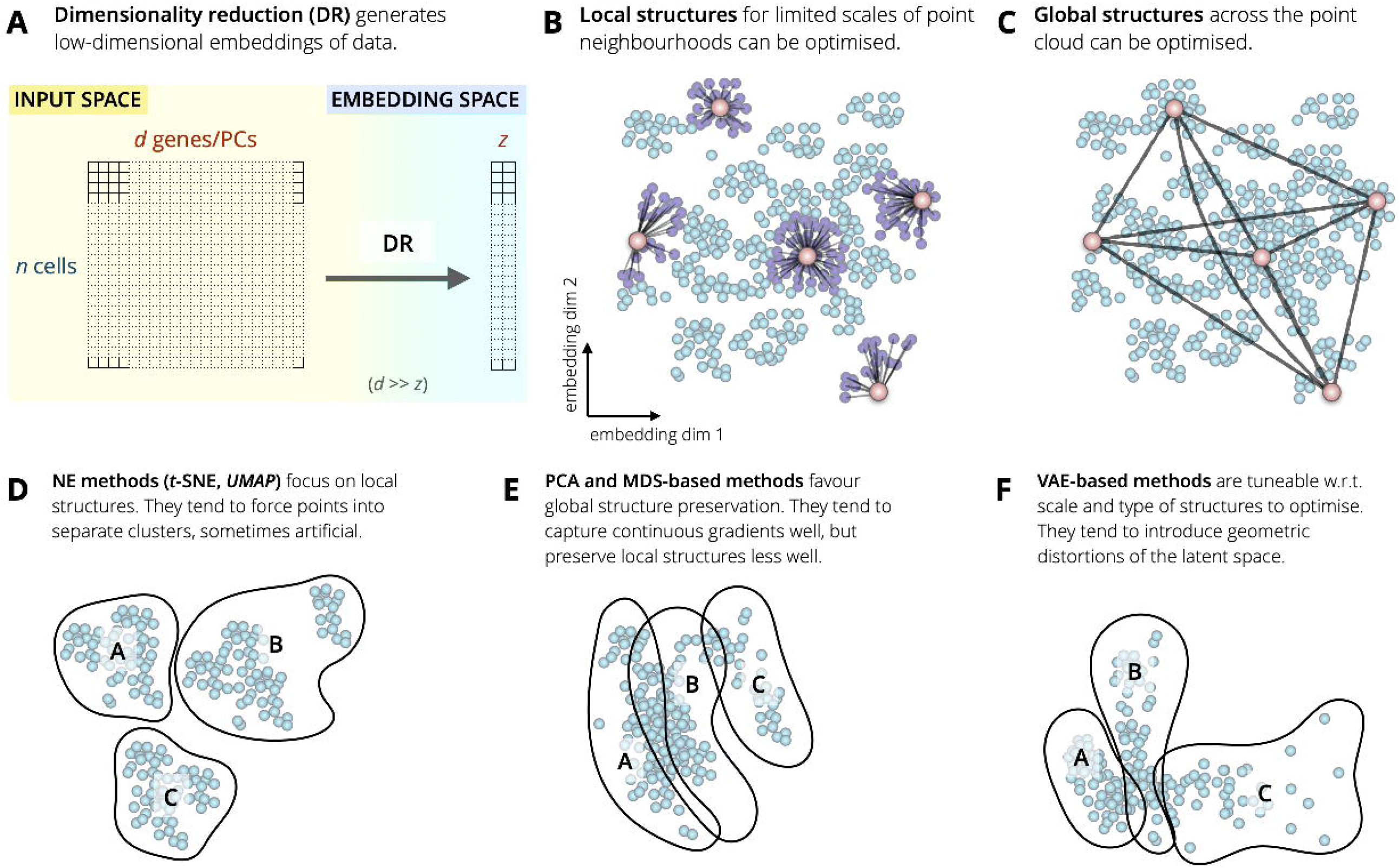
Schematic overview of dimensionality reduction and structure preservation. **(A)** Tabular *d* -dimensional input data is transformed into a *z*-dimensional embedding by DR. While the original *d* features correspond to specific genes or principal components (PCs), the *z* features of the embedding do not. The number of data points *n* does not change between the inputs and the embedding. **(B, C)** DR can focus on preserving neighbourhood relationships between points, respectively, at a localised scale or globally. **(D)** Neighbour embedding (NE) methods are typically ‘cluster-forming’. **(E)** Principal component analysis (PCA) and methods based on multidimensional scaling (MDS) do not have a locality bias and favour global (multiscale) structure preservation. **(F)** Methods based on variational autoencoders (VAEs) are highly flexible in the objectives they optimise. The models’ embedding spaces tend to introduce geometric distortions with respect to the input data.

Current methods differ considerably in the criteria they optimise. *Principal component analysis* (PCA) is a popular deterministic method that obtains ordered orthogonal vectors explaining the most variance in the data: *principal components* (PCs). To reduce dimensionality, a set number of top PCs is taken as the basis for a LD vector space, used for projection. PCA is widely applied to preprocess HD sparse data, including scRNA-seq^6^. It is typically used to generate 50-to 100-dimensional representations amenable to downstream processing, including further DR. Alternatively, the top 2 or 3 PCs can be taken directly as the embedding. However, this inherently limits the ability to capture complex non-linear relationships between features, as PCA only uses linear transformations.

Non-linear DR methods overcome this, with *t* -SNE^7^ and *UMAP* ^8^ having gained special prominence in single-cell biology^2,9^. They both fall into the family of *neighbour embedding* (NE) algorithms, and can be formulated as optimising point positions via attractive and/or repulsive forces^10^. NE algorithms aim to preserve local point neighbourhoods, and are suitable for visualising clusters and comparing alternative labellings. However, their inherent *‘locality bias’* can distort the relative positions of cohesive cell clusters, introducing artificial gaps and arbitrary global layouts^11^. By *locality bias*, we mean that local structures in the data (bounded by some *‘scale’* parameter that governs size of the point neighbourhoods of interest), are optimised (see Figure 1**B**). While *scale* can be increased (*e*.*g*., the *perplexity* parameter in *t* -SNE or neighbourhood count in *UMAP*), the limitation to local structures remains. Some initialisation strategies partially compensate for this, yielding better preservation of large-scale structures^2^.

Current single-cell bioinformatics literature lacks formal definitions of the terms ‘local’ and ‘global’ that would lend themselves clearly to extensive comparative evaluation across many datasets. There has been an emphasis on evaluation metrics that are straightforward to use in benchmarks involving representative data^12–14^. This has been in lieu of extensive descriptions of the underlying data manifolds, their intrinsic dimensionality, and dataset-specific topologies, which is infeasible with much of single-cell omics. The present study, similarly, focuses on broadly applicable strategies that make few assumptions about the dataset in hand.

Our work here, while focused on application more than on theory, offers a conceptual framework for the local/global dichotomy. By ‘local structures’ we refer to structures in some close neighbourhood of any given vantage point in the input data, with neighbourhood size governed by the scale parameter described above. Conversely, global structures are not limited by a scale parameter (Figure 1**C**) and can extend further across the point cloud under this conceptual framework. The term ‘structure preservation’ (SP) is then used to indicate a measure of how well the original HD structures are preserved in the LD embeddings (despite inherent constraints due to reduced dimensionality).

A number of recent methods have set out to increase the preservation of global structures, or to find a favourable local-global balance. Some, including *TriMap* ^15^ and *PaCMAP* ^16^, are variants of NE. While NE methods are typically susceptible to partitioning data points into artificial ‘islands’ (Figure 1**D**), both *TriMap* and *PaCMAP* sample some points outside of local neighbourhoods in the optimisation process to improve the global layout of embeddings.

*SQuad-MDS* ^17^ is a recent adaptation of *multidimensional scaling* (MDS), which preserves relative HD pairwise distances between some points regardless of neighbourhood or structure scales. In practice, this optimises the global layout of embeddings, similarly to PCA despite algorithmic differences (Figure 1**E**).

Models such as *scvis* ^18^ and *ivis* ^12^ are examples of variational autoencoders (VAEs;^19^), a flexible neural network architecture. VAEs embed HD data in a smooth latent space, favourable for representing continuous transitions between cell states (Figure 1**F**).

Overall, the trend of attempting to better balance local and global SP can reduce misinterpretations of *t* -SNE and *UMAP*, which are often applied to data with important global or hierarchical structures, despite not being optimised to represent them faithfully^20,21^. However, a major open problem remains. This is the issue of systematically evaluating the accuracy of embeddings. Recent comparisons apply both unsupervised and supervised scores^13,14,22^, often via measuring performance in downstream inference tasks, such as clustering or classification^12,23^. These approaches exhibit some shortcomings. First, when scoring local structure preservation (SP), setting a hard threshold for a ‘local’ neighbourhood^13^ is potentially heavy-handed. Second, to assess global SP some metrics effectively downsample data, computing distances between representative points determined via clustering^13^, thus sacrificing rigour. Third, evaluation of embeddings via downstream tasks^14^ yields a proxy score, fundamentally dependent on the classification or clustering algorithm used.

More broadly, a recent critique^24^ questioned the usefulness of single-cell DR due to failures to preserve distances between points. A response to it^25^ demonstrated that *t* -SNE and *UMAP* preserve point neighbourhoods and separate biologically relevant clusters. This further underscores a need for clear and fair evaluation criteria and indicators of embedding trustworthiness.

Our *ViScore* framework addresses the concerns regarding evaluation and builds on the conceptual ‘local-versus-global’ distinction we drew above. *ViScore* introduces highly scalable unsupervised SP scoring with few assumptions on the data and embeddings (novel *R*_NX_ curve approximations), an improved qualitative supervised evaluation method (*Extended Neighbourhood-Proportion-Error* ; xNPE), and a fully-fledged benchmarking framework, all of which we describe in this paper.

With an empirical evaluation framework in place, we set out to develop a DR model for representing both local and global structures faithfully. *ViVAE*, our flexible deep learning-based method, exhibits a favourable balance between the two criteria. Furthermore, it incorporates *encoder indicatrices* (EIs): a tool for visualising unwanted distortions in existing embeddings. Grounded in previous work within differential geometry^26^, EIs serve as a straightforward QC check that is model- and dataset-specific and allows the investigator to pinpoint specific parts of the embedding that are artificially expanded, shrunk, or stretched. The aim of EIs is to boost the adoption of quality-control and explainability measures in DR, which has been limited so far. They are a flexible approach that represents as much information about embedding distortions in a 2-dimensional projections as possible, while remaining easy to interpret.

The following *Results* section describes our methodological contributions and validates them in an empirical study. First, we present the *ViScore* approach to SP scoring. Second, we employ it in a quantitative comparison of *ViVAE* to other DR methods. Third, we perform a case study with a longitudinal developmental dataset^4^ to validate the capacity for multi-scale SP by *ViVAE* in an applied context with known trajectory structures. Fourth, we demonstrate the use of EIs as a tool for on-demand detection of distortions, using the same dataset. Finally, we compare meaningful differences between *t* -SNE, *UMAP*, and *ViVAE* on a non-developmental dataset^27^, evaluating localised distortions with the use of more focused, supervised evaluation measures also implemented in *ViScore*.

### Concepts of ‘local’ and ‘global’ across disciplines and applications

The notions of local and global scale are highly pertinent to the presented work. We note that understanding of these terms differs across disciplines of study.

In topology and graph theory, ‘local’ and ‘global’ are complementary scales of description: the local focuses on what happens in a small neighbourhood, while the global asks how those parts fit into an entire system. In topology, the distinction is central to the difference between neighbourhood-level geometric information and whole-space invariants such as connected components or holes^28^. In graph theory, it appears in the contrast between measures based on a node’s immediate neighbours, such as degree (*i*.*e*., number of connections), and measures that depend on paths or structure across the entire graph, such as centrality^29^.

In applied topology and topological data analysis (TDA), local methods are used to capture fine-scale geometric variation, noise structure, and neighbourhood connectivity in data. In graphs and networks, local structure usually means the direct neighbourhood of a node: adjacent node counts, how tightly those neighbours connect, and how clustered the surrounding subgraph is.

A global perspective, on the other hand, asks about properties of the entire space that cannot be recovered from one neighbourhood alone. In topology, classic invariants such as connected components and loops are global because they describe the overall shape of the object rather than one local region. In graphs and networks, global measures depend on relationships spanning much or all of the graph, so they reflect a node’s position in network-wide flow, connectivity, or integration. Graph-theoretic work on local-global convergence makes this distinction precise by separating descriptions that depend only on bounded neighbourhoods from those that incorporate larger-scale organization^30^.

In single-cell bioinformatics, ‘local’ and ‘global’ are used in multiple contexts. Some previous studies have applied notions from topological data analysis to describe local and global scales in the context of gene expression data^31,32^. However, within the subfield of dimensionality reduction (DR) and visualisation, the distinction in scale is often used in a highly operational way, with less focus on theoretical underpinnings. Here, local structure broadly refers to preserving relative neighbourhoods of similar cells, whereas global structure means preserving broader relationships among more distant cell populations, such as developmental branches, trajectories, or inter-cluster separation^13,14^.

The terms usually refer to the faithfulnesss of an embedding or manifold representation with respect to the original input data, not directly to topological invariants or network metrics. By contrast, in topology the local/global distinction is about the mathematical scale at which a property is defined, and in graph theory it is about whether a measure depends on immediate neighbourhoods or on the graph as a whole.

## RESULTS

First, we present a framework for evaluating DR in both supervised and unsupervised settings (with and without the use of cell labels, respectively). This is implemented in the *ViScore* Python package, which includes an automated large-scale benchmarking solution.

Figure 2**A** introduces *R*_NX_ curves^33^ as the basis for assessing *structure preservation* (SP). This is a mathematical framework for quantifying preservation of structure at any given scale. Crucially, we solve the outstanding issue of poor scalability of *R*_NX_ with a custom approximation algorithm that extends their use to large scRNA-seq data. This has been unfeasible previously, thus dramatically restricting the evaluation of single-cell DR. We use *R*_NX_ to derive two metrics which score, respectively, local (locally biased) and global (scale-agnostic) SP, and use them to compare *ViVAE* to other methods.

**Figure 2.**
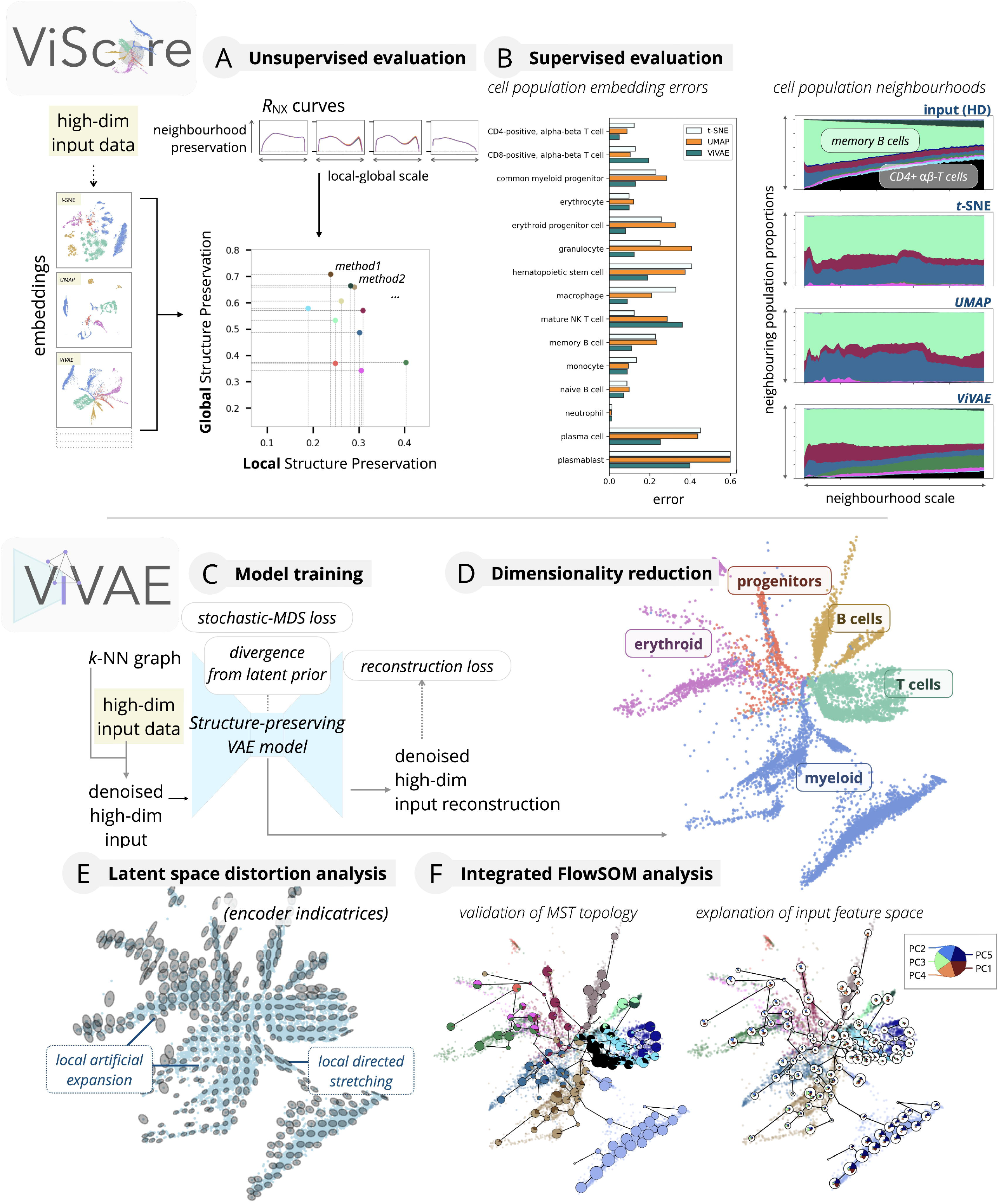
Schematic overview of proposed workflows. **(A)** Unsupervised evaluation based on *R*_NX_ curve analysis quantifies neighbourhood preservation for local and global structures. **(B)** *(Left side*.) Supervised evaluation via *Extended Neighbourhood-Proportion-Error* (xNPE) quantifies embedding error per cell population. *(Right side*.*) Neighbourhood composition plots* (NCPs) reveal inaccuracies in relative positions of cell populations in evaluated embeddings. **(C)** Input data is denoised and used to train a regularised VAE model. **(D)** The trained model is used to produce a low-dimensional embedding for visualisation. **(E)** Encoder indicatrices (EIs) evaluate distortions of the latent space learned by the model’s encoder. **(F)** The model can be used to validate a *FlowSOM* tree topology *(left side)* and project input feature values onto representative clusters *(right side)*.

Figure 2**B** (left side) shows the *Extended Neighbourhood-Proportion-Error* (xNPE): a proposed score for supervised (cell label-based) evaluation. xNPE, based on the NPE^22^, measures embedding quality per cell population. It captures distortions that change the relative positions between labelled cell populations. Providing cell population-level scores instead of embedding-level ones, xNPE serves as a QC check especially for comparing multiple candidate embeddings of the same dataset.

Another QC tool shown in Figure 2**B** (right side) are *neighbourhood composition plots* (NCPs), which diagnose sources of mis-embedding. They provide a qualitative description of errors, in contrast to the quantitative approach of the xNPE. NCPs visualise, for a given population of interest, which cells surround it, respectively, in the HD input space and in the evaluated embedding. This is done using a range of local neighbourhood scales.

Second, with a suitable evaluation framework in place we developed *ViVAE* : a DR model for balanced local and global structure preservation. This is a regularised variational autoencoder (VAE) that learns non-linear transformations of scRNA-seq data (Figure 2**C**). The input data is first denoised to prevent the model fitting to noise. Next, the model is trained on the denoised data with the use of a proposed stochastic multidimensional scaling (*stochastic-MDS*) regulariser. Stochastic-MDS is a fundamental extension of the ‘stochastic quartet’ approach^34^, making adaptations to yield a differentiable loss function that is usable in a deep learning context. In practice, stochastic-MDS improves the preservation of global structures and embedding distortions (as shown further).

The trained model can generate data embeddings (Figure 2**D**) and map previously unseen data points from the same domain as training data, facilitating the assessment of technical artifacts by comparing data batches.

Figure 2**E** introduces *encoder indicatrices* (EIs): a QC tool that allows the investigator to visually inspect unwanted distortions which the model introduces into the embedding. Appearing as ellipses in the figure, EIs denote both the direction and the magnitude of unwanted expansion, shrinkage, or stretching of parts of the embedding space, to the extent that these can be visualised in 2-*d*.

Additionally, *ViVAE* integrates with *FlowSOM* ^35^, which couples clustering and graph embedding via minimum spanning trees (MSTs), as shown in Figure 2**F** (left side). This helps in validating the MST topology against the embedding, and vice versa, and examining whether expected ontologies, hierarchies, or developmental lineages are reflected. Projecting values from the input feature space onto representative clusters also quickly captures major sources of variability, as a dataset-specific explanatory mechanism (right side).

Both *ViScore* and *ViVAE* are available in dedicated online repositories, with several guided tutorials, experiments, and benchmarking guidelines provided alongside the software itself.

### *ViScore* evaluates structure preservation across multiple scales

The proposed *ViScore* methodology assesses the preservation of high-dimensional (HD) structures in their low-dimensional (LD) embeddings across different scales. The term *‘Structure Preservation’* (SP) is taken to denote the degree to which an embedding retains the neighbourhood structures between HD points in the original data. The choice to focus on Euclidean distance-induced neighbourhoods, rather than distances themselves, is motivated by a previous analysis demonstrating their conceptual relevance within single-cell DR^25^.

Figure 3**A** shows an example of 4 alternative embeddings of a developmental dataset^4^. A single ‘vantage-point’ cell is chosen (orange cross); a colour gradient shows the true neighbourhood-ranks of all other cells with respect to the vantage point. These neighbourhood-ranks are computed from the HD input data, not the embeddings.

**Figure 3.**
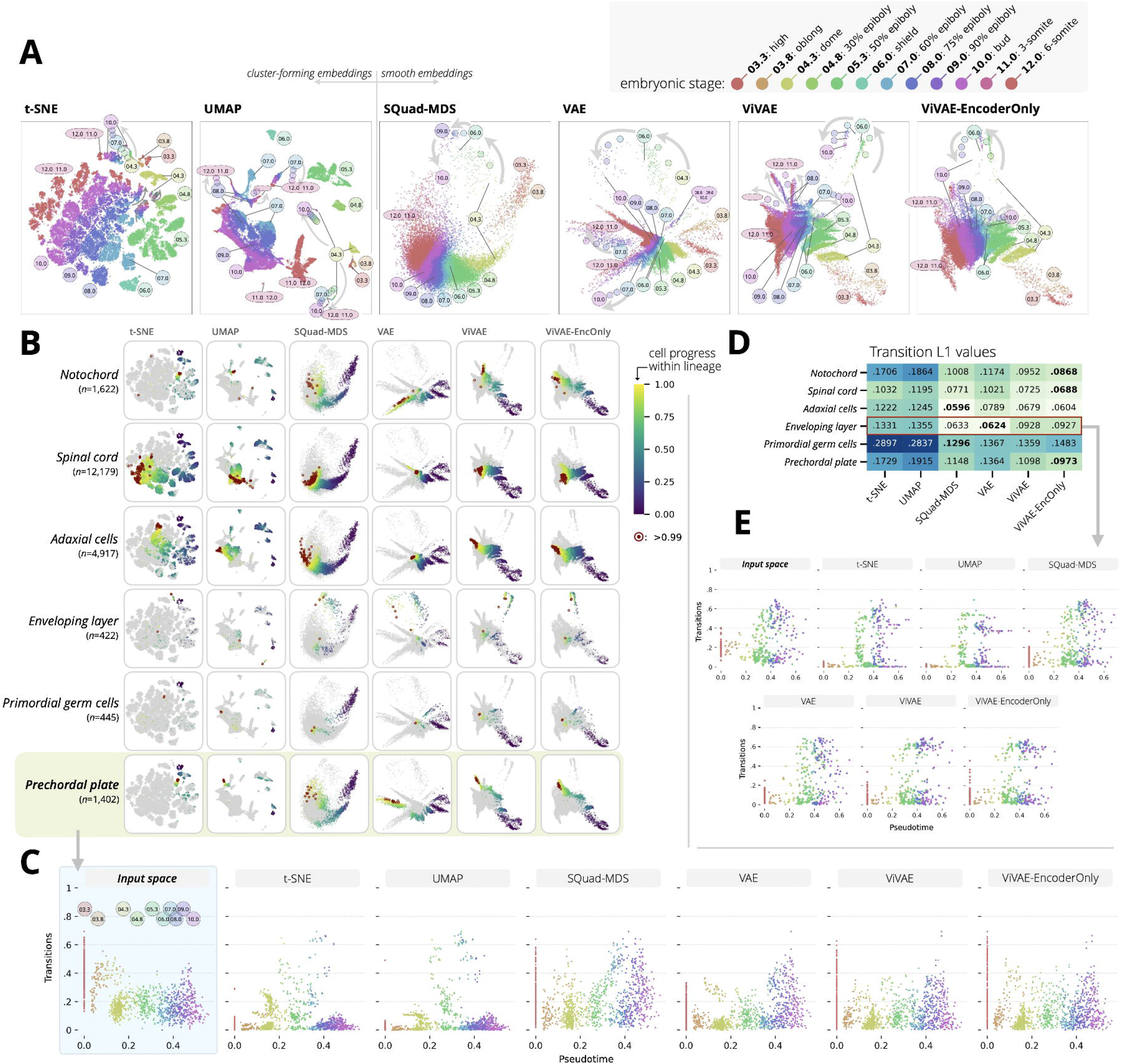
*Farrell* dataset embeddings with developmental lineages in zebrafish embryos. **(A)** 2-d embeddings of *Farrell* data^4^ by *t* -SNE, *UMAP, SQuad-MDS*, VAE, *ViVAE*, and *ViVAE-EncoderOnly* are shown, with colour-coding by embryonic stage. The layout of cell subsets by stage is indicated in each embedding, showing major developmental gradients using *hpf* labels and arrows indicating progression. **(B)** Six annotated developmental lineages are projected onto each embedding, using colouring by pseudotime-derived rank, with terminal cells in the highest one percent highlighted in red. **(C)** Scaled distances between cells along the *prechordal plate* lineage are shown for input data and the embeddings. **(D)** Average L1 distances between inputspace and embedding distances for lineages shown in *B*. are plotted in a heatmap. **(E)** Transition distances for the enveloping layer lineage validate that trajectories start diverging around the 4.3- or 4.8-*hpf* stage.

*t* -SNE and *UMAP* separate the data into islands, achieving a high level of detail (note that these separations always need to be validated as non-artificial). On the other hand, the gradient of neighbour ranks is inconsistent: this demonstrates failure to preserve some structures. *SQuad-MDS*, at the other extreme, shows minimal separation into islands, but the gradient is preserved better. Finally, *ViVAE* balances these two criteria (as examined further in the following section).

Despite under-defined notions of ‘local’ and ‘global’ in relevant literature, we devise exact definitions for a *Global SP* score and *Local SP* score, both based on *R*_NX_ curves^33^ (see *Methods*). This is shown in Figure 3**B**. It is crucial that we avoid setting a hard neighbourhood-size cut-off for what constitutes a ‘local’ structure, and we compute Global SP as a truly scale-agnostic score (*i*.*e*., small-scale and large-scale structures are treated as equally important), thereby reducing arbitrary designs.

In overview, both SP scores measure the preservation across all possible scales; for any point in a dataset of size *N*, these are scales from 1 to *N* −1. While *Global SP* treats all scales as equally important for the final score (*scale-agnostic*), *Local SP* applies a *log*_10_ rescaling of neighbourhood scales, which upweights the importance of smaller-scale structures (*locally biased*, as proposed in an earlier study^33^).

The SP scores are upper-bounded by 1, with 0 corresponding to a random embedding (baseline), and negative values to a worse-than-random representation of neighbourhood structures. A realistic upper bound, however, is dataset-specific, since full preservation of neighbourhoods would only be possible for some cases of noiseless data lying on a 2-manifold that can be flattened. Determining this upper bound would translate to describing the true manifold of each single-cell dataset; this is not currently feasible. However, based on comparative analyses presented here and elsewhere^13,14^, we know some DR methods to effectively prioritise Global SP (PCA, *SQuad-MDS*) while others (mainly *t* -SNE) focus on Local SP. This knowledge serves as an empirically determined anchor for good performance with respect to each criterion.

### *ViVAE* is a parametric model that balances local and global structure

We compare 12 DR models run on 8 publicly available scRNA-seq datasets with respect to Local and Global SP. These models are PCA, *UMAP* ^8^, *DensMAP* (a *UMAP* variant), *t* -SNE^7^, *PHATE* ^23^, PaCMAP^16^, TriMAP^15^, SQuad-MDS^34^, ivis^12^, VAE (a standard variational autoencoder) and our own: *ViVAE* (default) and *ViVAE-EncoderOnly*. While default *ViVAE* is trained using reconstruction loss, a variational regulariser, and stochastic-MDS, *ViVAE-EncoderOnly* omits reconstruction loss (and does not have a decoder). This yields a loss function for the *ViVAE* and *ViVAE-EncoderOnly*, respectively:

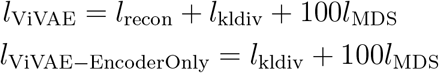

where *l*_recon_ is the autoencoder reconstruction loss (mean squared error), *l*_kldiv_ is the Kullback-Leibler divergence from an isotropic latent prior standard to VAEs and *l*_MDS_ is the proposed stochastic-MDS loss. *Figure S5* displays the values of *l*_recon_, *l*_kldiv_, and *l*_MDS_ during training per epoch, comparing a vanilla VAE model, *ViVAE*, and *ViVAE-EncoderOnly* using 2 training datasets. We elaborate on the used models, their hyperparameters, and the comparative analysis set-up in *Methods*.

We test the DR methods on the following datasets:

- *Qiu* ^36^: 89*k* T cells from mouse embryos. Chosen as immunologically relevant data from a common model organism.
- *Reed* ^27^: 25*k* immune cells from human breast. Chosen due to containing widely studied immune cell compartments, and because DR methods struggle to separate some of the labelled cell populations clearly (*e*.*g*., NK cells within the T+NK compartment, which we evaluate in a corresponding case study). Also as a representative of non-developmental data, for which cluster-forming embeddings (*t* -SNE, *UMAP*) are traditionally used.
- *Suo* ^37^: 908*k* human pre-natal T and NK cells across organs. Chosen as a recent representative of a large multi-organ dataset with developmental gradients.
- *Strati* ^38^: 33*k* human bone marrow cells from healthy donors. Chosen due to the prominence of bone marrow lymphopoiesis as use case in trajectory inference, as well as its relevance in immunology and haematology.
- *TabulaMuris* (subset)^39^: 40*k* mouse bone marrow cells. Chosen as a popular mouse dataset complementary to *Strati*.
- *TabulaSapiens* (subset)^40^: 22*k* human thymic cells. Similarly to *Strati*, chosen due to the prominence of T-lymphopoiesis as a use case in TI analysis and its relevance in immunology and hematology.
- *Shekhar* ^41^: 45*k* cells from mouse retina. Chosen as a popular transcriptomic dataset where a documented batch effect can be revealed by non-linear DR.
- *Farrell* ^4^: 39*k* cells from zebrafish embryos at different developmental stages. Chosen to evaluate the preservation of smooth gradients and discontinuities, both of which are present, due to a longitudinal component. The provided annotation of developmental lineages allowed us to validate embeddings of developmental pathways in a dedicated case study.

Following a standard preprocessing protocol^2,12^, we normalise and scale expression matrices from each dataset and apply PCA to extract orthogonal components as input features, retaining 100 PCs (as a more conservative approach). The full preprocessing workflow is given in *Methods* and the online *ViScore* repository.

Figure 3**C** compares Local and Global SP by dataset, averaged across 5 repeated runs. For all datasets, at least one of the *ViVAE* models lies on the Pareto front, meaning no other model outperforms it at both tasks. Notably, *t* -SNE gives the highest Local SP wherever applied (training timed out at 4 hours for the largest *Suo* dataset) and *SQuad-MDS* gives the highest Global SP across all datasets. *TriMap*, on the Pareto front in 6 out of 8 experiments, achieves similar Local SP to *UMAP*, while improving Global SP, which aligns generally with previously published results^14^.

*Table S1* contains results with and without the use of denoising of input data for each method, which amounts to an ablation study that demonstrates a boost to Local SP with denoising with VAE-based models. Additionally, the *ViVAE* and VAE models in our comparison use the same architecture, therefore comparing them yields an ablation study that demonstrates a boost to Global SP with stochastic-MDS. *Table S2* gives *p*-values from one-sided Wilcoxon tests comparing the default and encoder-only variant of *ViVAE* with each other method per dataset for Local SP, Global SP, and a tentative ‘Balanced SP’ metric (computed as the harmonic mean between the two, as described in *Methods*). The Local, Global, and Balanced SP means and standard deviations are plotted separately in *Figure S1*. The change to *R*_NX_ curves induced by input data denoising is shown in *Figure S2*. Additionally, *Figure S3* shows the impact of varying target dimensionality (from 2 to 3, 5, and 10 dimensions) on structure-preservation capacity of different models using the *Farrell* dataset, with higher latent dimensionality generally yielding higher SP. *Figure S4* shows the effects on SP of varying the ‘scale’ parameter for those methods that have one, again with the *Farrell* dataset.

While these SP results give an overview of different methods’ capabilities to represent data faithfully, there are other notable aspects of *ViVAE*. Based on a VAE architecture, *ViVAE* is a parametric model with a latent space that is amenable to QC and explainability measures. This includes encoder indicatrices (as demonstrated below) and integration with *FlowSOM* (demonstrated in online materials). Furthermore, *ViVAE* does not have an effective ‘scale’ hyperparameter determining the size of relevant point neighbourhoods for which structures are optimised during training, unlike *t* -SNE, *UMAP, DensMAP, PaCMAP, TriMap, PHATE*, or *ivis*. This approach, using the truly multi-scale stochastic-MDS loss, simplifies potential hyperparameter tuning.

In the following sections, we demonstrate ways in which *ViVAE* meaningfully outperforms its alternatives in two case studies: one with the developmental *Farrell* data and one with the non-developmental *Reed* data. The *Farrell* case study shows clearly why the local-global SP balance achieved by *ViVAE* is advantageous.

### Stochastic-MDS regularisation in *ViVAE* achieves more faithful embeddings of developing cell lineages in zebrafish embryos

The *Farrell* Drop-Seq dataset^4^ aggregates 38,731 cells from 12 annotated developmental stages across 694 zebrafish embryos, from 3.3 to 12 hours post-fertilisation (*hpf*). In addition to *hpf* - sorted embryonic stage labels, the authors provide annotations of developmental lineages and inferred pseudotime values inducing the ordering of cells within each lineage. As shown in our comparison results, the ‘*ViVAE*’ and ‘*ViVAE-EncoderOnly*’ models achieve high Global SP, lower only than *SQuad-MDS*, which they exceed at Local SP. Both *ViVAE* models improve Local and Global SP over a standard VAE.

Figure 4**A** compares embeddings from *t* -SNE and *UMAP* (examples of neighbour-embedding; NE), *SQuad-MDS* (example of multidimensional scaling; MDS), and VAE, default *ViVAE*, and *ViVAE-EncoderOnly* (examples of variational autoencoders; VAEs). Embryonic stages are marked by colour and schematic annotation bubbles with arrows in the direction of development (increasing cell differentiation). As a generally known phenomenon, the NE methods separate data into clusters, with the rest forming smoother embeddings. Of the smooth embeddings, *SQuad-MDS* shows the least separation of 11- and 12-*hpf* cells into distinct branches, with VAE or default *ViVAE* separating them the most; panel **B** maps these separations onto trajectory endpoints. The smooth embeddings place most 6-*hpf* (‘shield phase’) cells clearly along the major trajectory, in contrast to NE embedding separating them out. The correctness of the smoother embeddings is supported by lower Extended Neighbourhood-Proportion-Error (xNPE) embedding errors for shield phase cells: ranging from 0.781 in *t* -SNE, 0.858 in *UMAP* and 0.620 in *SQuad-MDS* to 0.243 in VAE, 0.242 in *ViVAE*, and 0.128 in *ViVAE-EncoderOnly*. Lower xNPE indicates a more accurate layout of this population with respect to others (see *Methods*).

**Figure 4.**
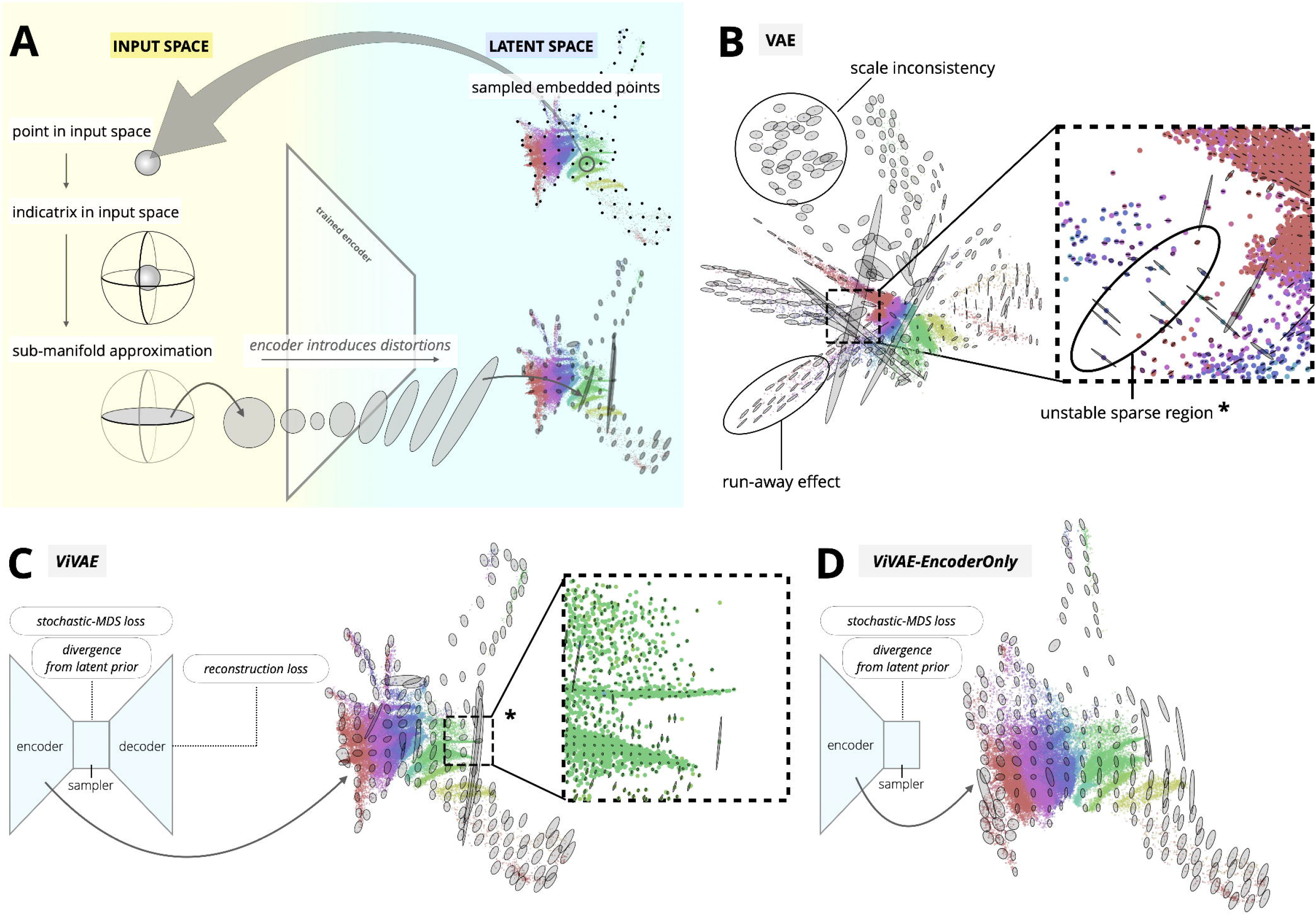
Analysis of latent space distortion in VAE-based embeddings of the *Farrell* dataset. **(A)** Overview of *encoder indicatrix* (EI) computation, using an embedding of the *Farrell* dataset^4^. Points in input space are obtained via grid-sampling from the embedding. Using local linearisations of the trained encoder, HD indicatrices for the points are flattened along corresponding horizontal tangent spaces, yielding a piecewise approximation of the embedding 2-manifold. Flattened indicatrices are transformed by the encoder and are plotted over the embedding. **(B)** EIs reveal commonly observed distortions of VAE latent space: scale inconsistencies, run-away effect inducing directed stretching and instabilities in sparse regions. **(C)** The default *ViVAE* model gives less varied indicatrices but some instabilities in sparse regions remain. **(D)** The additionally evaluated encoder-only *ViVAE* model learns a latent space that exhibits gradual distortion of scale and shape and less drastic instabilities due to sparsity (less stretching and fewer changes in direction of the stretching across different parts of the embedding space).

Figure 4**B** shows 6 distinctive developmental lineages, with cells coloured by normalised ranks along the progression. Terminal cells in each lineage are highlighted in red. Of particular interest is the challenge of embedding the *enveloping layer cell* (ELC) and *primordial germ cell* (PGC) lineages, terminating at 10.0-*hpf*. They contain few cells and are distinct from other lineages, as described in the original study. All 6 embeddings reflect this, with both lineages partially separating from others. Additionally, both lineages show some degree of bifurcation. The pathways are highly fragmented in *t* -SNE and *UMAP*, while in the smooth embeddings they retain more continuity, despite the low cell counts.

This raises the question of whether smooth embeddings represent transitional states correctly. Consider that this data is longitudinal, still allowing for gaps in trajectories between *hpf* stages. To test this, we quantify how continuous or fragmented each lineage is in the input (HD) space. In Figure 4**C**, we show Euclidean distances between cells ordered by pseudotime (‘transitions’) within the *prechordal plate* lineage. This is computed for input space and for each of the embeddings, and transformed to account for scale differences (via *min-max* and *log1p* scaling). Large distance values at the 3.3- and 3.8-*hpf* stages in input space validate that the less dense representation of these cell subsets by smooth embeddings is correct; *t* -SNE and *UMAP* collapse them artificially.

To measure the distortion of these transitions by each embedding, we report L1 distances between the input and embedded distance distributions, normalised by each lineage’s total cell count. These values (Figure 4**D**) show that the smooth DR models indeed achieve a more faithful representation of the lineages. Furthermore, Figure 4**E** shows transition distances in the ELC lineage to validate the bifurcation observed in embeddings. The real separation is shown to be gradual, more similar to the representation by the smooth embeddings.

### Encoder indicatrices reveal distortions in VAE model space

We propose *encoder indicatrices* (EIs): a tool to visualise distortions of (V)AE-based latent spaces, and use them to analyse VAE, default *ViVAE*, and *ViVAE-EncoderOnly* embeddings of the *Farrell* data. Figure 5**A** gives an overview of the algorithm. In each case, an embedding is generated by a trained encoder model. Representative points are sampled from the embedding using a uniform grid. They are then mapped to their coordinates in the input (HD) space. For each sampled point, a small hypersphere around it forms its HD *indicatrix*. Here, ‘indicatrix’ denotes a shape that helps us describe how the particular part of the point cloud gets projected by the trained model. Using a local linearisation of the encoding, we then flatten the indicatrix and transform it using the encoder again.

**Figure 5.**
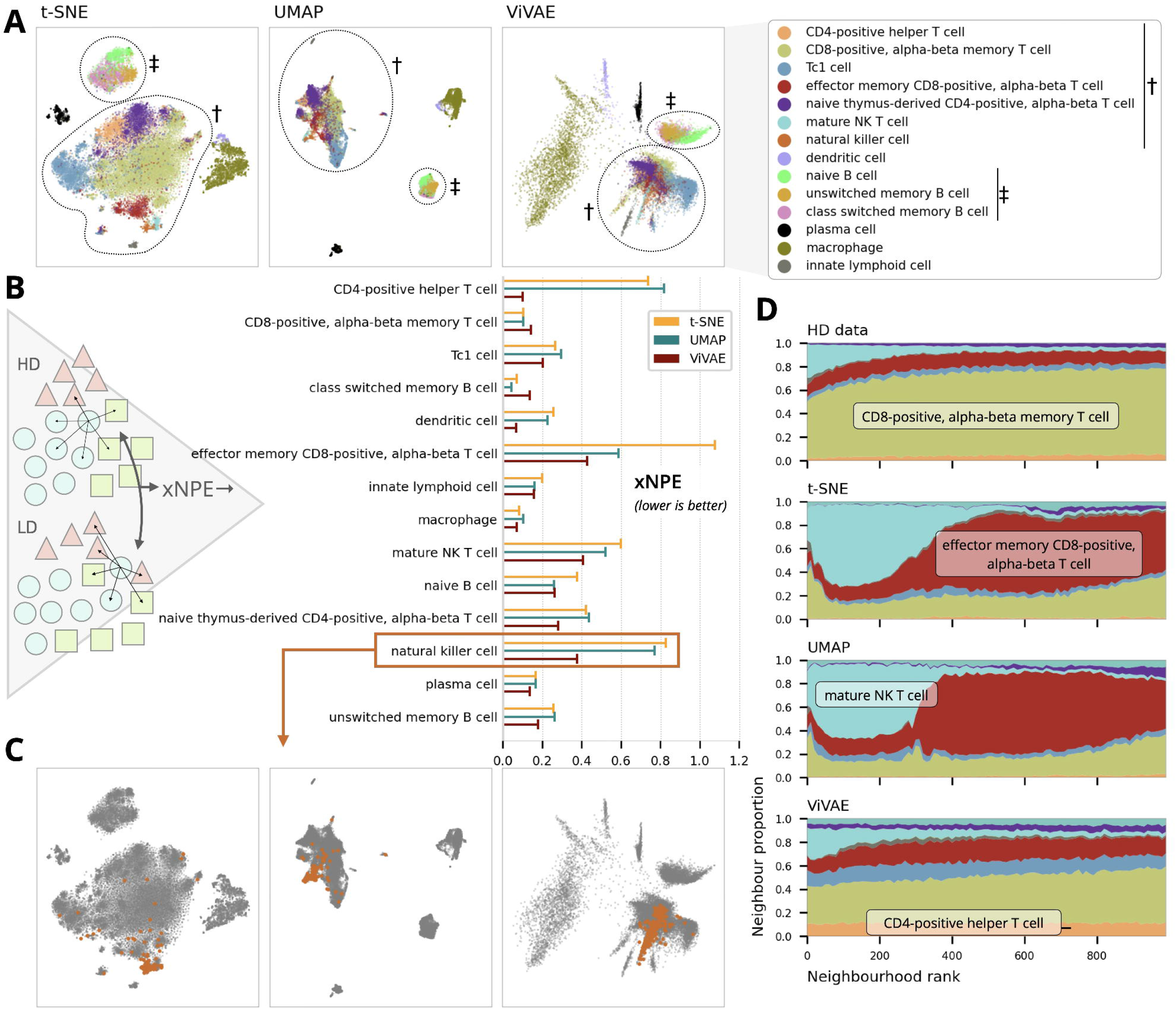
*Reed* dataset embedding and analysis of population-wise error. **(A)** 2-d embeddings of the *Reed* data^27^ from *t* -SNE, *UMAP*, and *ViVAE* are shown, with populations indicated by colour, according to legend on the right. Symbols mark the the T- and NK-cell (†) and B-cell (‡) compartments. **(B)** xNPE (population-wise embedding error) is computed using 1000-nearest-neighbour relations in input space and embeddings. Distributions of neighbour identities in terms of self and non-self are compared. **(C)** NK cells are highlighted in orange as population-of-interest in each embedding. **(D)** Neighbourhood composition plots (*k* = 1000) show distribution of other populations in the neighbourhoods of the population-of-interest. Colour-coding from panel *A* is used, along with text annotations.

This transformation introduces size and shape distortions, which get mapped onto the embedding coordinates. In this way, we objectively describe local embedding artifacts throughout the latent space, to the extent that this is achievable in a 2-dimensional visualisation. In line with previous related work^26^, we adopt the use of ellipses superimposed on the embedding for a readily interpretable visual description of artifacts.

Figure 5**B** uses EIs to highlight commonly observed embedding distortions in VAEs. First, scale inconsistences make some parts of the latent space artificially sparser. Second, a ‘runaway’ effect causes directed stretching of some point subsets. Third, some sparse regions are ill-defined, showing drastic extension where the embedding should be smoother.

Figure 5**C** shows that the default *ViVAE* model, with denoising and stochastic-MDS regularisation, gives more uniform indicatrices, reducing scale inconsistencies and run-away effects. Some instabilities in sparse regions as seen with VAEs remain. Figure 5**D** shows that the encoder-only *ViVAE* model creates a latent space with some gradual scale and shape distortion, but further reduces instabilities due to sparsity. Overall, this shows a notable reduction of latent space distortion in *ViVAE* over a standard VAE.

### *ViVAE* achieves structure-preserving reduction of non-developmental immune cell data dimensionality

Here we demonstrate the usefulness of *ViVAE* as a general-purpose DR algorithm for other than developmental data, where Global Structure Preservation (SP) by smoother embedding algorithms is less of a focus. To that end, we demonstrate novel supervised (label-based) evaluation tools from *ViScore*: the Extended Neighbourhood-Proportion-Error (xNPE) and neighbourhood composition plots (NCPs).

Specifically, we use the *Reed* immune cell dataset, originating from a single-cell transcriptomic atlas of the adult human breast^27^. As opposed to the *Farrell* embryological data (in previous section), we expect more discontinuity between cell types, as the breast tissue is seeded with distinct populations of mature immune cells, likely presenting a challenge to our smoother VAE-based embedding method.

Figure 6**A** compares *ViVAE* to the prevalent NE methods (*t* -SNE and *UMAP*), showing a degree of major cell-compartment separation in each. Since DR invariably introduces artifacts, we seek to quantify and describe them here, using a higher-resolution approach than embedding-level SP scoring. Figure 6**B** quantifies population-level embedding errors using xNPE, which leverages the provided cell labelling. For each *population of interest* (POI), the xNPE aggregates the distribution of ‘self versus non-self’ neighbours (how many neighbouring cells belong to the POI itself, versus a different population) over all the cells in the POI. It does so separately in the HD input space and each LD embedding, ending up with two distributions. These two distributions are then compared using the earth mover’s distance (EMD). The lower the EMD value, the lower the distribution of the POI is with respect to the ground truth.

By this metric, *t* -SNE fares less well than *UMAP* for 8 of 14 populations and less well than *ViVAE* for 12 of 14 populations, despite a high Local SP score based in an unsupervised setting. Importantly, in contrast to Local SP, xNPE only focuses on how each population interfaces with the others around it; small positional inaccuracies or swapping positions within the population are not penalised. (The online *ViScore* repository includes a dedicated proof-of-concept experiment with synthetic data.)

We investigate how the natural killer (NK) cells population is distributed in different embeddings, picking them as the POI because all three embeddings place them within the same compartment as T cells (marked ‘†’ in panel **A**), but assign them somewhat scattered positions within it (orange points in panel **C**). Both *UMAP* and *ViVAE* outperform *t* -SNE in terms of xNPE. To derive a more meaningful interpretation, we generate neighbourhood composition plots (NCPs) for the NK-cell population (Figure 6**D**). These are stacked area plots showing smoothed proportions of neighbouring populations across different neighbourhood sizes (up to 1000), using a sliding-window approach. These proportions can be compared between input (high-dimensional, non-reduced) data and the different embdeddings, and they exclude neighbouring cells from the POI itself.

According to the top NCP, the NK cell population is close to a CD8^+^ - T-cell population (olive green) in input space. All three embeddings underestimate this (the population is underrepresented), but *ViVAE* does so the least. Notably, *t* -SNE and *UMAP* show an intrusion of mature NK T cells (light blue) into a close neighbourhood of the NK cells, meaning their proximity is exaggerated; *ViVAE* avoids this artifact. The proximity to CD4^+^ helper T cells (orange), relative to the HD expression data, is artificially reduced by *t* -SNE and *UMAP* and increased in *ViVAE. t* -SNE and *UMAP* also mis-embed NK cells with respect to effector memory CD8^+^ - T cells, in contrast to *ViVAE*.

This case study shows separation of major immune cell compartments by *ViVAE* and low population embedding errors, validating its competitiveness at general-purpose single-cell data embedding, and thus extending its functionality beyond visualising trajectory data. The supervised evaluation metrics presented here can leverage known cell population labelling, making it possible for the domain expert to examine embeddings of single-cell data more meaningfully, in an ad-hoc setting.

## DISCUSSION

We have shown *ViVAE* to achieve favourable multi-scale structure preservation and faithful embedding of both trajectory and non-trajectory data, according to comparisons with other methods on an array of real biological datasets as well as focused case studies. Integrating denoising of inputs and a novel stochastic-MDS component with a VAE model, *ViVAE* balances the continuous nature of VAE latent spaces with an ability to separate major cell compartments. To improve interpretability, we equip *ViVAE* with QC functionality in the form of *encoder indicatrices*; crucially, these can be implemented for any differentiable DR model. In addition to a regularised VAE, we propose and evaluate a decoder-less model (*‘ViVAE-EncoderOnly’*). We take the full model as default due to mostly slightly better preservation of local structures, but evaluate both.

We have demonstrated that balancing local and global structure preservation can lead to more faithful embeddings of developmental, longitudinal, but also non-trajectory (single-snapshot) data. In essence, the proposed methodology is designed for a broad range of exploratory data analysis workflows.

Throughout the work, we use Euclidean distances in PCA-reduced input space, which yield evaluations that are generally congruent with the known trade-offs between local and global structure preservation by established methods. However, for adaptability to sparser and higher-dimensional data (*e*.*g*., expression of a filtered gene set, without initial PCA reduction), we implement the use of cosine distances as an alternative for use with the stochastic-MDS loss, as well as in the unsupervised structure-preservation scoring.

The development of *ViVAE* followed our initial implementation of model-agnostic structure preservation scores in *ViScore*, which were necessary for a quantitative evaluation. Overcoming previous conceptual and computational limitations, we coin unsupervised embedding-level scores that quantify structure preservation without using hard local/global cut-offs, landmark points based on clustering, or other previously proposed proxy measures. In addition, we introduce supervised population-level evaluation metrics for comparing embeddings and explaining artifacts.

Our contributions to evaluation and interpretability are motivated by increasing sizes of single-cell datasets, the availability of large public databases and atlases^42–44^, and ongoing efforts to systematically evaluate single-cell data analysis algorithms^11^. DR currently aids in exploration and hypothesis formulation, and its usage will likely rise with more public data. This will require efforts to prevent erroneous conclusions from spurious patterns or artifacts in embeddings.

Having adapted notions from differential geometry^26^ to visualise distortions in the model space using *encoder indicatrices*, we estimate that an increased use of such explainability tools will help in describing method-specific artifacts. Future endeavours in interpretable non-linear single-cell DR may involve hierarchical embeddings^45^ and interactive tools for evaluating embedding quality at local levels^46,47^.

### Limitations of this study

Another major avenue of research involves topological data analysis (TDA) to describe point clouds. We acknowledge that concepts such as local and global structure have undergone thorough and formal examination within this field^28^. Further, some studies have sought to tackle the problem of interesting structures existing at multiple scales in single-cell data^48,49^. Nonetheless, this work within TDA has not been fully incorporated within the single-cell DR field. While interesting work on topology-preserving autoencoders exists^50^, our familiarity with a recent study on topological regularisation of autoencoders for single-cell DR^51^ emphasises to us that dataset-specific descriptions of topologies can prove cumbersome and non-trivial.

More broadly, describing the true manifolds of real biological data is currently intractable. Therefore, the focus of our study here, and a handful of others that compare DR methodologies^12–14,25^, is instead on empirical evaluation across different datasets that are representative of a range of real-world scenarios.

In line with this approach, our work seeks to provide ways to measure and optimise multiscale structure preservation without requiring too much tuning or too many assumptions on underlying structures. This is why our current study does not use TDA and remains restricted to the use of nearest-neighbour graphs in lieu of more elaborate topologies.

Nonetheless, our ongoing research is focusing on alternative graph structures for both denoising and DR evaluation, such as networks from the Louvain algorithm^52^, witness complexes^49^, or branched spanning trees robust to noise^53^. Our integration of *ViVAE* with *FlowSOM* ^54^ (shown in Figure 1) is a tentative step toward the use of graphs together with embedded point clouds to capture relevant structures.

## RESOURCE AVAILABILITY

### Lead contact

Requests for further information and resources should be directed to and will be fulfilled by the lead contact, David Novak.

### Materials availability

The proposed software packages are implemented in Python and available in dedicated GitHub repositories: *github*.*com/saeyslab/ViVAE* and *github*.*com/saeyslab/ViScore*

### Data and code availability

This paper analyzes existing, publicly available data. Sources of datasets are listed in the key resources table (KRT).

All original code has been deposited at GitHub and is publicly available as of the date of publication. The dedicated repositories are listed below.

- The *ViVAE* software package and code required to reproduce all case studies in this paper are publicly available in the *saeyslab/ViVAE* GitHub repository as of the date of publication.
- The *ViScore* software package and code required to reproduce the quantitative comparison of *ViVAE* and other dimensionality reduction methods are publicly available in the *saeyslab/ViScore* GitHub repository as of the date of publication.

Any additional information required to reanalyze the data reported in this paper is available from the lead contact upon request.

## ACKNOWLEDGMENTS

D.N. was funded by the FWO (Fonds Wetenschappelijk Onderzoek) and this research was conducted as part of the FWO Strategic Basic research project 1S40421N. C.d.B. was a beneficiary of an FSR Incoming Post-doctoral Fellowship from UCLouvain during part of this work. P.L. is a FRIA grantee with the Belgian FNRS (Fonds National de la Recherche Scientifique). J.A.L. is a Research Director with the Belgian F.R.S.-FNRS (Fonds National de la Recherche Scientifique). S.V.G. is supported by an FWO postdoctoral research grant (1272823N, Research Foundation – Flanders). This work was supported by the Flemish Government under the Flanders AI Research Program (Project 174K02325).

We thank Sebastian Damrich for his invaluable advice regarding the implementation of encoder indicatrices and Anna Konstorum for her advice and guidance regarding supervised evaluation that allowed us to build on top of the Neighborhood Proportion Error algorithm.

## AUTHOR CONTRIBUTIONS

D.N. conceived of and implemented the *ViVAE* methodology, supervised evaluation metrics in *ViScore*, and the large-scale evaluation framework, designed and ran experiments, and drafted the manuscript. C.d.B. conceived of and implemented the *R*_NX_ curve approximation and wrote the corresponding manuscript section. C.d.B., P.L., J.A.L., S.V.G., and Y.S. provided extensive feedback on the manuscript. Y.S. edited the manuscript. S.V.G. and Y.S. supervised the study.

## DECLARATION OF INTERESTS

The authors declare no competing interests.

## SUPPLEMENTAL INFORMATION INDEX

**Document S1**. PDF file with supplemental figures and notes. **Figure S1**: visualization of structure-preservation results as means and standard deviations per score. **Figure S2**: visualization of R_NX_ curves per dataset and method. **Figure S3**: visualization of the effects of varying target dimensionality on the structure-preservation (SP) scores. **Figure S4**: visualization of the effects of varying the ‘scale’ hyperparameter on the structure-preservation (SP) scores. **Figure S5**: values of different loss function terms during training for a VAE, *ViVAE*, and *ViVAE-EncoderOnly* model. **Note S1**: pseudocode for the *Denoising* algorithm. **Note S2**: pseudocode for the *Encoder Indicatrices* algorithm. **Note S3**: pseudocode for the *Extended Neighbourhood-Proportion-Error* algorithm. **Note S4**: Python code for scRNA-seq data preprocessing. **Note S5**: key resources table (KRT)

**Table S1**. Full results of structure-preservation comparative analysis of dimensionality reduction methods

**Table S2**. *p*-values from one-sided Wilcoxon tests for the *ViVAE* and *ViVAE-EncoderOnly* models, corresponding to results in *Table S1*

**Table S3**. Specification of dimensionality reduction methods, their Python implementations, and hyperparameter values used for training

## STAR METHODS

### Key resources table

Key resources table (KRT) is provided as *Note S5*.

### Method details

#### Variational autoencoders for dimensionality reduction

An autoencoder (AE) is a type of neural network trained to reconstruct an input **X**_**n**×**d**_ ∈ 𝒳 as an approximation 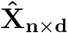 while compressing **X** into a lower-dimensional representation **Z**_**n**×**z**_ ∈ 𝒳 (where 𝒳= ℝ^*d*^, 𝒳= ℝ^*z*^, *z < d*)^19^. It consists of two stacked feed-forward networks: the encoder *E*_Φ_ : 𝒳 →𝒵, which transforms **X** to **Z**, and the decoder *D*_*θ*_ : 𝒳 →𝒳, which transforms **Z** to 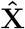. The network parameters (encoder weights Φ and decoder weights Θ) are learned so as to reduce a reconstruction loss: in *ViVAE*, input data is scaled and reconstruction loss is computed as the mean square error 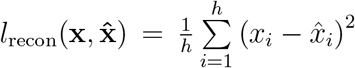 for **x** and 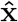 being points or batches of points drawn, respectively, from **X** and 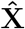. Each node of the network transforms inputs using a non-linear activation function, allowing for learning complex, non-linear patterns through back-propagation of error at output.

The variational autoencoder (VAE) is a generative model based on the AE. Instead of encoding and decoding a latent representation **Z** directly, a distribution 𝒟 in latent space is learned, and a sampler generates a non-deterministic representation **Z** and decodes it. The dissimilarity (Kullback-Leibler divergence) between 𝒟 and a latent prior distribution (typically isotropic Gaussian) is used as an additional loss term *l*_kldiv_ during training. The model is trained in mini-batches drawn from **X** to minimise *l*_recon_ and *l*_kldiv_ jointly.

Since the VAE learns a salient representation **Z** of the original data **X**, we can use **Z** for visualisation of important high-dimensional structures in **X** that are initially obscured. This gives us a parametric DR model. By default, our *ViVAE* model uses the Gaussian error linear unit (GELU) activation function and a symmetric architecture of 32,64,128 32, and 2 (*z*) nodes, respectively, in consecutive encoder layers, and a reverse of this for the decoder. The model generalises to higher latent space dimensionalities. The default optimiser used in training is Adam, with a learning rate of 0.001 and weight decay of 0.0001.

#### PCA-initialisation using imitation loss

In our online *ViVAE* repository, we implement an optional pre-training procedure for PCA initialisation of (V)AEs. We devise the *imitation loss*, which computes L2 distances between embedded points and the first two PCs of the input dataset. In practice, pre-training the model to imitate PCA prior to the full training stabilises results, reducing variability between repeated runs of the training.

#### Input denoising

The denoising algorithm used here is based on an approach described in^55^ and applied previously in^49^; it is equivalent to the mean shift algorithm^56^. The procedure is written out using pseudocode in *Note S1*. Denoising requires a *k*-nearest-neighbour graph (*k*-NNG) over preprocessed input data; to this end, we use a fast approximate *k*-NNG construction algoritm^57^. By default, we take *k* = 100. Each point is then moved closer toward the average coordinates of its *k* nearest neighbours, where a hyperparameter *λ* determines size of the shift. This can be done in multiple iterations, each shift potentially exposing the underlying manifold and making the point cloud less scattered. In our workflow, we use *λ* = 1 and apply 1 iteration of denoising to input data. One may experiment with these parameters, or apply denoising to LD embeddings (using the *k*-NNG in input space) to achieve a sharper defined shape of embedded cell clusters or pathways.

According to ablations conducted in our full comparative analysis (results in *Table S1*), the regularised VAE model used in *ViVAE* is particularly amenable to input denoising, achieving better Local SP in the resulting embedding. Crucially, we still use the original (preprocessed, but non-denoised) input data as ground truth in evaluating the embeddings, so as to ensure a fair comparison between set-ups. We test all of the evaluated methods with and without denoising, further concluding that the structure-preserving properties of *ViVAE* are not exclusively attributable to the denoising step.

#### Stochastic-MDS loss and encoder-only model

Similarly to some other VAE-based DR algorithms (*scvis* ^18^, *ivis* ^12^, *VAE-SNE* ^58^), *ViVAE* formulates constraints on the LD embedding with respect to HD inputs. We achieve this using a *stochastic multidimensional scaling* (*stochastic-MDS*) loss function. This is an adaptation of the stochastic quartet loss introduced in *SQuad-MDS* ^17,34^, which we previously generalised and reimplemented as *group loss* for use in differentiable models in a preliminary study^59^.

For stochastic-MDS, samples in each training mini-batch drawn from **X** are divided randomly into quartets (group of 4 points) and a cost value is computed based on the difference between normalised distances within the quartet in **X** and within the corresponding embedded quartet in **Z**. An update to each sample’s position in LD is computed by differentiating a cost function with respect to all of the 6 intra-quartet distances. Therefore, each sample’s position in LD moves to jointly minimise the error of its quartet as a whole. For data points indexed by *i* and *j*, we use pairwise Euclidean distance functions, *δ*_*ij*_ in HD and *d*_*ij*_ in LD (an option of using cosine distances is also given). We compute relative distances between pairs of points among quartets as

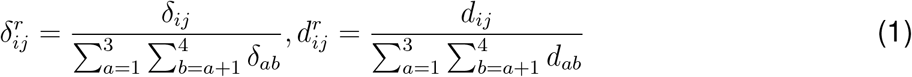

The loss function is then defined as

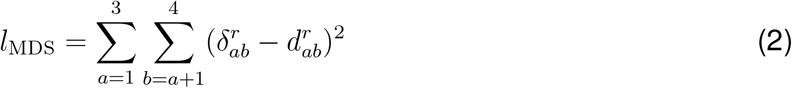

To increase coverage of different parts of the point cloud, we re-sample quartets multiple times (4 times by default) and average out the computed gradients at each pass. Regarding our fixed choice of 4 points as group size, our past experiments^59^ showed no improvement in terms of structure preservation when group size was increased.

For our default ‘*ViVAE*’ model (labelled that way in our comparative analysis results), we use a weight factor of 1 for *l*_recon_, 1 for *l*_kldiv_ and 100 for *l*_MDS_: this amounts to a combination of the structure-preserving capabilities of a VAE and the MDS model. We also evaluate a ‘*ViVAE-EncoderOnly*’ model, which assigns a weight of 0 to *l*_recon_, 1 to *l*_kldiv_ and 100 to *l*_MDS_. In this case the decoder is not used, and we obtain a probabilistic MDS model consisting only of an encoder and a sampler. Technically, this is then not an autoencoder, but we use the name *ViVAE* because other parameters of the model are kept fixed between this model and the default one, yielding an ablation experiment for the use of reconstruction error.

Regarding our choice of loss term weights, we observe and report good performance with these default weights across the datasets included in our comparison, at least as long as a consistent data preprocessing workflow is used. We ensure that the scale of the stochastic-MDS loss term is invariant with respect to chosen batch size and number of quartet re-samplings. Our software implementation enables the usage of cosine distances in HD for computing the stochastic-MDS loss. This is explored in the online *ViVAE* repository.

#### Encoder indicatrices for detection of local distortion

During training, AEs and VAEs both learn a smooth latent space, defined by the encoder. The parametric nature of these models, along with the fact that the transformations they learn are differentiable, allow us to explore properties of this latent space. Specifically, we can detect local contraction, expansion and directed stretching of its different parts.

Our methodology extends the work on geometric regularisation for (non-variational) AEs^26^. To visualise undesirable artifacts, the use of *indicatrices* ^60^ is proposed to quantify latent space distortion. An indicatrix can be understood, in a simplified way, as a circle on the surface of a small hypersphere in the original (HD) ambient space, embedded into the (LD) latent space. In a previously published study^26^, multiple indicatrices from different parts of the HD point cloud are sampled and then pulled back into the latent space through the decoder, to reveal distortions introduced by the model.

A perfect, non-distorted embedding results in all indicatrices remaining circular and identical (circular because shape is preserved and identical because there are no scale inconsistencies across different parts of the latent space). The distortion that each indicatrix reveals is specific with respect to the ambient space, transformation function and the coordinates of the sampled point. In the previous study we reference^26^, indicatrices are computed for the transformation learned by the decoder: we can refer to them therefore as ‘decoder indicatrices’. Crucially, decoder indicatrices quantify distortion of latent space specifically with respect to the decoded reconstruction 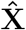.

In contrast, we derive a solution to the ‘encoder indicatrix’ problem, *i*.*e*., quantifying local embedding (and latent space) distortion with respect to the original high-dimensional inputs. Since a general analytical solution is not feasible (essentially due to the encoder not being injective), we provide a numerical solution, described in *Note S2*. A schematic overview of the algorithm is given in Figure 5**A**.

To compute encoder indicatrices, we begin by sampling points from the existing embedding closest to uniform grid over it and mapping them onto the respective high-dimensional inputs *X*_*s*_. For each point *x* ∈ *X*_*s*_, we compute the Jacobian matrix 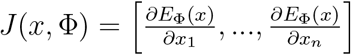 which, when developed, has dimensions *z* × *d* (*z* being latent dimension and *d* being input dimension), and gives a local linearisation of the encoder with respect to this point. Under the assumption that the *E*_Φ_ function is approximately linear for very small neighbourhoods of any given *x* ∈ *X*_*s*_, only *z* vectors from a small hypersphere around it are needed. This is because all other orthogonal vectors will be sent to zero by *E*_Φ_.

We obtain the *z* useful vectors as the bases of the *horizontal tangent space* specific to the given point and the encoder. The horizontal tangent space is approximated using singular value decomposition of the Jacobian matrix. We then sample points from a small circle in this approximation of the horizontal tangent space for the queried point. The sampled points are taken as the flattened indicatrix, which is then transformed by the encoder to reveal the local size and shape distortions with respect to input data. Same as in the decoder indicatrix case, non-distortion yields identical, circular indicatrices. In practice, the indicatrices differ in size and are stretched into ovals. Note that the use of indicatrices with a linear AE would result in a degenerate case^26^.

#### Integration with *FlowSOM*

To boost interpretability of *ViVAE* embeddings for practitioners familiar with *FlowSOM*, we provide the option to superimpose the topology of a *FlowSOM* minimum spanning tree (MST) along with median marker expression, proportion of manually labelled population, or metacluster assignment of clusters over the *ViVAE* embedding. This is demonstrated in our online *ViVAE* repository. In practice, we adapt code from the recent Python clone of *FlowSOM* ^54^ to enable the projection of clusters into the latent space of a trained model.

#### Rank-based neighbourhood preservation scoring

We base our unsupervised quality-assessment measures on widely adopted rank-based DR evaluation criteria^61^. The basis for these is the concept of a *co-ranking matrix*, which quantifies the changes of ranks of neighbours to each point in terms of pairwise distances in LD, as compared to HD.

For any pair of HD points, we denote distance from *ϵ*_*i*_ to *ϵ*_*j*_ as *δ*_*ij*_. Conversely, we denote the distance from an LD point **x**_*i*_ to **x**_*j*_ as *d*_*ij*_. We define the rank in HD of *ϵ*_*j*_ with respect to *ϵ*_*i*_ as the set cardinality *ρ*_*ij*_ = ∣*k* : *δ*_*ik*_ *< δ*_*ij*_ ∨ (*δ*_*ik*_ = *δ*_*ij*_ *∧ k < j*) . (This can be re-stated more simply as ‘*ϵ*_*j*_ is the *ρ*_*ij*_-th neighbour to *ϵ*_*i*_’.) The corresponding rank in LD is similarly defined as *r*_*ij*_ = ∣{*k* : *d*_*ik*_ *< d*_*ij*_ ∨ (*d*_*ik*_ = *d*_*ij*_ ∧ *k < j*)}. The non-reflexive (non-self) neighbourhoods of *ϵ*_*I*_ and **x**_*i*_ are then written as 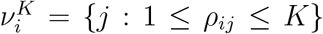 and 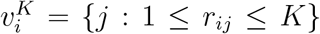 respectively, for a neighbourhood size of *K*. We then define the co-ranking matrix for a dataset of size *N* as **Q** = [*q*_*kl*_]_1*≤k,l≤N−*1_, where *q*_*kl*_ = ∣ (*i, j*) : *ρ*_*ij*_ = *k* ∧ *r*_*ij*_ = *l*}∣. Any off-diagonal positive values in this matrix indicate a change of rank in the LD embedding versus the HD input dataset, and are undesirable. The co-ranking matrix records changes in the order of relative distances between points as we go from the original, high-dimensional data, toward a lower-dimensional embedding. A reduction in rank is labelled an *intrusion* and is recorded left of the diagonal, whereas an increase is termed an *extrusion*, and is recorded right of the diagonal. A change in rank that crosses a given close-neighbourhood upper bound of *K* (a sample enters or leaves the *K*-ary neighbourhood of its reference) is termed a *hard-K intrusion* or *hard-K extrusion*, respectively. Based on^33^, we measure structure preservation at different scales by recording the changes in rank within a limited neighbourhood size *K* as we expand size from 1 to *N* -1. We start with defining the average structure preservation in a *K*-ary neighbourhood as

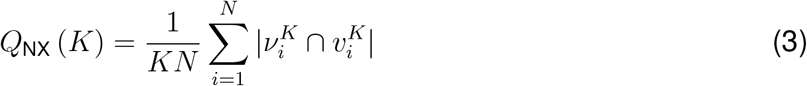

and record these values for the different values of *K*. We denote this as the *Q*_NX_ curve. A rescaling of the *Q*_NX_ curve is the *R*_NX_ curve; it shows improvement of neighbourhood preservation over a random embedding, which would result in an average 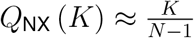. This is given by

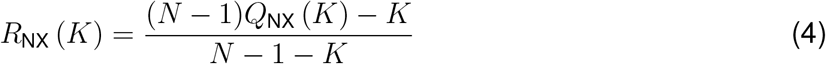

for 1 ≤ *K* ≤ *N* − 2. The *R*_NX_ curve helps evaluate how the structure of the HD dataset is preserved across different scales in the LD embedding.

#### Multi-scale evalution of structure preservation

In reference to concepts introduced in the previous section, we denote the area under the *R*_NX_ curve with a linear (non-transformed) scaling of *K* as the global structure-preservation score (*s*_*g*_, or ‘Global SP’). We also report the area under the *R*_NX_ curve with a logarithmic (log_2_) scale for *K*, up-weighting local point neighbourhoods, as the local structure-preservation score (*s*_*l*_, or ‘Local SP’). The Global SP score is computed as the area-under-curve

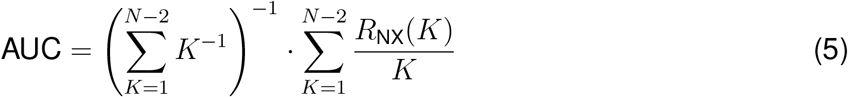

while the Local SP uses *log*_2_*K* instead of *K* (otherwise the same equation is used). Additionally, we compute a balanced structure-preservation score (‘Balanced SP’) *s*_*b*_, calculated as the harmonic mean (*F* -measure) between the *s*_*l*_ and *s*_*g*_:

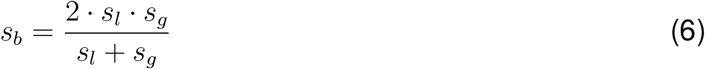

However, we recommend looking at the values of *s*_*l*_ and *s*_*g*_ separately when possible, since overwhelming performance in only one of them can still result in high *s*_*b*_ in cases where this is unintuitive.

It is noteworthy that evaluating *Q*_NX_ and *R*_NX_ curves for *K* ranging from 1 to *N* − 2 entails a 𝒪 (*N*^2^ log *N*) time complexity. Their computation requires sorting, for each data point, its HD and LD distances with respect to all *N* − 1 remaining data points^62^. These criteria, in their original implementation, are hence exhaustive and slow to assess the low-dimensional embeddings of very large datasets. We address this below by presenting a novel algorithm that circumvents scaling issues.

#### Q_NX_ and R_NX_ curve approximation

The 𝒪 (*N*^2^ log *N*) time complexity required to evaluate the unsupervised DR quality criteria for all neighbourhood sizes, reviewed above, is unaffordable in big data contexts. One could trivially display the quality curves for very small neighbourhood sizes only, smaller than 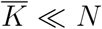^63^. The ensued time complexity would scale as 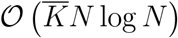, thanks to dedicated algorithms for fast nearest neighbour searches^57,64,65^. Nevertheless, such a trick cannot enable fairly comparing several DR methods, which may preserve structures and trends at different scales. Methods with global properties such as PCA would be systematically disadvantaged with respect to local schemes such as *t*-SNE. For this reason, quality curves should still be computed for a much wider proportion of neighbourhood sizes, to grasp the behaviour of DR methods at different data scales. We hence introduce an algorithm to approximate *Q*_NX_ and *R*_NX_ curves for large datasets quickly and with high precision, based on repeated sub-sampling via vantage-point trees.

One can first note that *Q*_NX_ gives a score computed with respect to each data point index *i* ∈ ℐ, with the set ℐ = {1, …, *N*} containing all point indices, averaged over the entire dataset. This score evaluates, for each data point, the proportion of its *K* nearest HD neighbours preserved in the LD embedding. Faithfully estimating this score average, however, may only require relying on a fraction of the data samples, instead of the whole dataset. In this view, a set 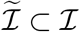 indexing data representatives could be defined to summarise the data cloud. A candidate approximation of *Q*_NX_ hence develops as

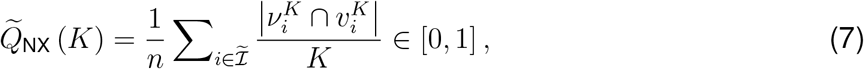

where 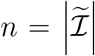. Compared to *Q*_NX_, the time complexity of evaluating 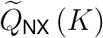 for all neighbour-hood sizes *K* ∈ {1, …, *N* − 1} drops to 𝒪 (*nN* log *N*); one must indeed only compute the *N* − 1 HD and LD distances with respect to each data representative indexed in 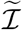, instead of all *N* data samples. From 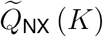, an approximation of *R*_NX_ (*K*) is obtained as

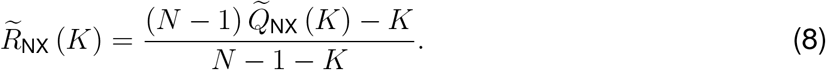

To define the set 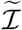, a first option consists in randomly sampling *n* representatives among the *N* data samples. Nevertheless, to ensure the consistency of the approximation of *Q*_NX_ (*K*) by 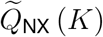, the samples indexed in 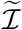 should faithfully represent the HD data distribution. On the other hand, the number *n* of representatives should remain small to keep the lowest possible time complexity. In this view, defining 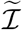 through pure random sampling indeed well summarises the HD data distribution, but only as soon as *n* gets sufficiently large, which undesirably increases running times. To keep the *n* value as small as possible while enforcing the representatives to cover the entire HD distribution, a well-balanced partition of the HD data distribution may first be computed. Space-partitioning trees such as vantage-point (VP) trees are extremely fast and efficient for this purpose^64,66^, being widely used to speed up nearest neighbour searches in big datasets^67,68^.

At each non-leaf node of a VP tree, a vantage point is first selected and the distances between the vantage point and the remaining data points are computed. The median distance is then computed in linear time, thanks to a quick-select procedure^69^. Two child nodes are finally created, respectively storing the remaining data points closer and further to the vantage point than the median. The median threshold ensures that the tree remains complete and balanced, with similar proportions of data points in all parts of the tree. Constructing the whole VP tree has (*N* log *N*) time complexity^64^. However, as only *n* representatives need to be determined, we may only grow the tree up to depth ⌊log_2_ *n*⌋, in 𝒪 (*N* log *n*) time. Indeed, the 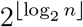leaves of the resulting tree define a balanced partition of the HD data, with each leaf containing a subset of the database. One can hence randomly sample a representative in each leaf, or possibly two if 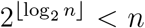, yielding *n* representatives which summarise the HD distribution well. Furthermore, only the leaves of the tree need to be stored, since only the latter are required to sample the representatives. The VP tree can thus be iteratively created in-place in a single array. In each node, the vantage point is selected as the furthest datum from the most central one.

Note that neighbour searches in VP trees do not scale well with the HD dimension due to the distance concentration^70^, which annihilates their time complexity advantage. Nevertheless, this only concerns neighbour searches performed in these trees, not their construction since in this case, data points are evenly spread with respect to the median distance to the vantage point. As we only seek to construct VP trees to partition the HD distribution without performing neighbour searches, they can be safely employed in our context without suffering from increasing HD dimension. As another advantage, VP trees solely require to be able to evaluate distances between the HD data points, without constraining them to be expressed as HD vectors. These trees can hence be generally computed even with complex data formats, unlike *k*-d trees for instance^66^, which involve Cartesian coordinates of the Euclidean space. For this reason, the size of *k*-d trees grows exponentially with the HD dimension, whereas the median threshold in VP trees prevents such increase, with depths scaling at most as 𝒪(log_2_ *N*).

Preliminary experiments revealed that *n* values between log *N* and 10 log *N* dramatically accelerate the DR quality criteria computation, while yielding close to perfect approximations of *Q*_NX_ and *R*_NX_; when considering big datasets with *N* growing to very large numbers, average scores estimated on small yet representative subsets of the database indeed intuitively quickly converge towards the actual average computed over the entire dataset. Values of ⌊10 log *N* ⌉ are hence subsequently employed for *n*, with ⌊·⌉ denoting rounding. Therefore, the time complexity required to evaluate 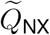 for all neighbourhood sizes scales as 𝒪 (*N* log^2^ *N*), which is considerably more affordable than the 𝒪(*N*^2^ log *N*) time required to evaluate *Q*_NX_.

#### Extended Neighbourhood-Proportion-Error

The *Neighbourhood Proportion Error* (NPE)^13,22^, quantifies preservation of the structure of manually defined populations of cells. We coin the *Extended Neighbourhood-Proportion-Error* (xNPE) as a modification of the NPE.

For each labelled population, NPE iterates over all the points it contains as vantage points. For each vantage point, the number of points in its *k*-ary neighbourhood belonging to the same population is counted. This is done for the HD input data, as well as the LD embeddings. The distribution of counts of same-population neighbours per vantange point is computed, and it is compared between HD and LD using total variation distance (TVD), resulting in a population-wise error score. The sum of these population-wise error scores is then returned as a measure of error for the entire embedding.

In xNPE, we use proportions of same-population neighbours rather than counts. Further, we use Earth mover’s distance (EMD)^71,72^ to compute differences between distributions, since it uses the entire distribution instead of the (more likely spurious) single largest difference between distributions, and return population-wise scores to avoid an assumption of consistent resolution of the cell population labelling.

Additionally, we scale each population-wise score. This is first done such that a score of 0 (best) is perfect and 1 (worst) is the upper bound which represents largest possible EMD value. By default, we then adjust the score again such that 1 corresponds to a random embedding baseline (as described in the corresponding pseudocode in *Note S3*).

Rather than summing or averaging population-wise errors to obtain an embedding-wise score, we inspect and compare the population errors to detect which parts of the input point cloud are the most mis-embedded. A proof-of-concept with synthetic data that compares xNPE to the unsupervised Local and Global SP scores is presented in the online *ViScore* repository.

#### Neighbourhood composition plots

To identify and visualise sources of embedding error at a population level, we use *neighbourhood composition plots* (NCPs). These plots visualise the composition of different-size neighbourhoods of a chosen labelled population of interest (POI) in HD data and its embeddings. This enables us to compare the position of the POI relative to other populations and how it is preserved in a given embedding.

For this, we query *k* nearest neighbours (*k* = 1000 by default) of each point in the POI, overlooking points in the POI itself. This is distinct from the xNPE approach described earlier, which makes the binary distinction between ‘self’ and ‘non-self’. While neighbourhood composition plots do not directly quantify introduced errors, they visualise the type of error (*e*.*g*., intrusion or extrusion of a specific population into or out of the neighbourhood of the POI). Composition of different-size (*K*-ary) neighbourhoods (1 to 100, with step size 10 by default) in terms of population labels is plotted for the POI as a stacked area chart, where each distribution of labels across the *K* axis is pooled for a range of neighbourhood ranks and for all points in the POI.

#### Model and hyperparameter specification

Model specifications and hyperparameters used in our quantitative comparisons of structure preservation (SP) by 12 DR methods, presented in *Results*, are given in *Table S3*.

The full configuration for our comparative analysis of structure preservation by different DR methods (as presented in *Results*) is available in the *ViScore* GitHub repository. It includes installation scripts for each method and instructions for deployment on a high-performance computing (HPC) cluster. We use default hyperparameters for running each method, except for larger batch size of 1024 for VAE-based models. For consistency, PCA-reduced data is used as input to each method, with reference to previously published standard preprocessing procedures^2,6^. While *PHATE* can be applied directly to the sparse count matrices from scRNA-seq experiments, it performs an initial PCA reduction anyway.

For ‘*VAé*, ‘*ViVAE*’, and ‘*ViVAE-EncoderOnly*’ we use 50 training epochs as default, because this training procedure is typically feasible on a consumer laptop within a few minutes even without the use of a dedicated GPU for acceleration. However, a higher number of training epochs typically leads to better convergence and structure preservation with many datasets. ‘*VAE*’ uses the same architecture and hyperparameters as ‘*ViVAE*’, so as to serve as a fair reference in an ablation experiment for the stochastic-MDS loss regularisation. Furthermore, for the comparison to simultaneously serve as ablation study for denoising, each method is evaluated both on non-denoised and denoised inputs. CPU-based methods (PCA, *t* -SNE (*scikit-learn* implementation), *UMAP* and *DensMAP* (*umap-learn* implementation), *PaCMAP, PHATE, TriMap*, and *SQuad-MDS*) were trained on a CPU cluster with an *Intel Xeon Gold 6140* processor. 16 GB of RAM were allocated for the processing of all datasets except *Suo*, for which 32 GB of RAM were allocated. GPU-based methods (VAE, *ivis, ViVAE*, and *ViVAE-EncoderOnly*) were trained on a GPU cluster with *Intel Xeon Gold 6242* and an *NVIDIA Volta V100* graphics card. 2 cores with 8 GB of GPU memory each were allocated.

#### scRNA-seq data preprocessing

Code and data to reproduce our study are openly available. We include instructions for downloading and preprocessing scRNA-seq datasets we used in our qualitative comparison in the *ViScore* GitHub repository, and in a Python code snippet in *Note S4*. The *scanpy* Python package (version 1.9.8)^73^ is used, and the preprocessing procedure is based on a previous study^2^, with the scaling step parametrised the same way as in the *ivis* workflow^12^.

Changes to this procedure or use of other data modalities may require corresponding changes to method hyperparameters, including individual loss term weights in *ViVAE*, for good performance.

### Quantification and statistical analysis

*Table S2* includes *p*-values from one-sided Wilcoxon tests comparing the default and encoderonly variant of *ViVAE* with each other method per dataset for Local SP, Global SP, and a tentative ‘Balanced SP’ metric. These were computed using the R programming language *wilcox*.*test* function.

### Additional resources

*ViVAE* repository: *github*.*com/saeyslab/ViVAE*

*ViScore* repository: *github*.*com/saeyslab/ViScore*

## Notes

https://github.com/saeyslab/ViVAE

https://github.com/saeyslab/ViScore

### Competing Interest Statement

The authors have declared no competing interest.

### Summary of Updates

Revised structure and format to conform to Cell Systems guidelines Incorporated context about notions of local and global in topology, graph theory

