## Supplementary material for "Interpretable models for scRNA-seq data embedding with multi-scale structure preservation": Document S1

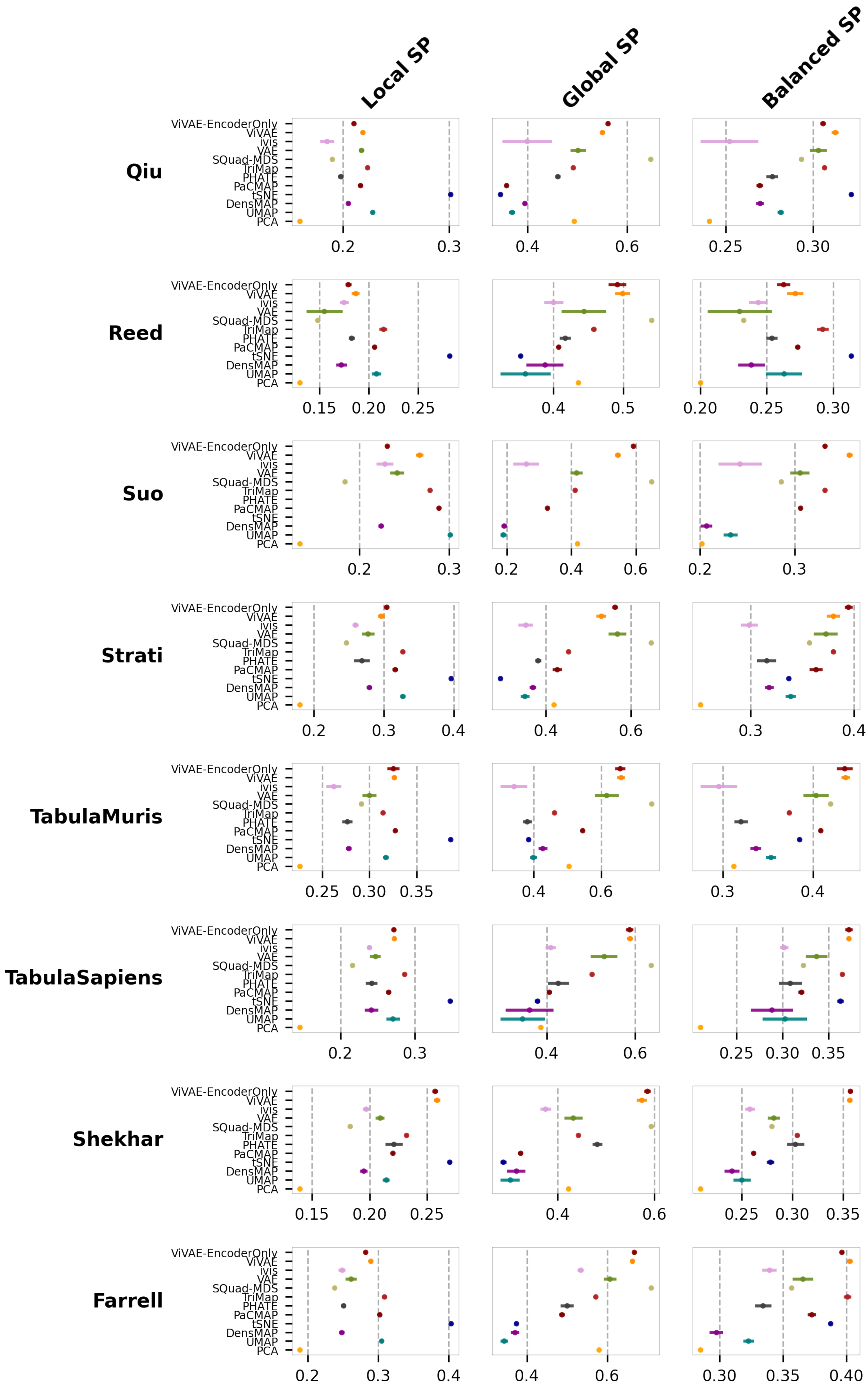


**Figure S1: Comparison of structure-preservation (SP) scores per dimensionality reduction (DR) method and dataset.** Results come from 5 runs with different random seed per method-dataset combination, with mean value (point) and 1 standard deviation (line) indicated. Local, Global, and Balanced SP are displayed separately.

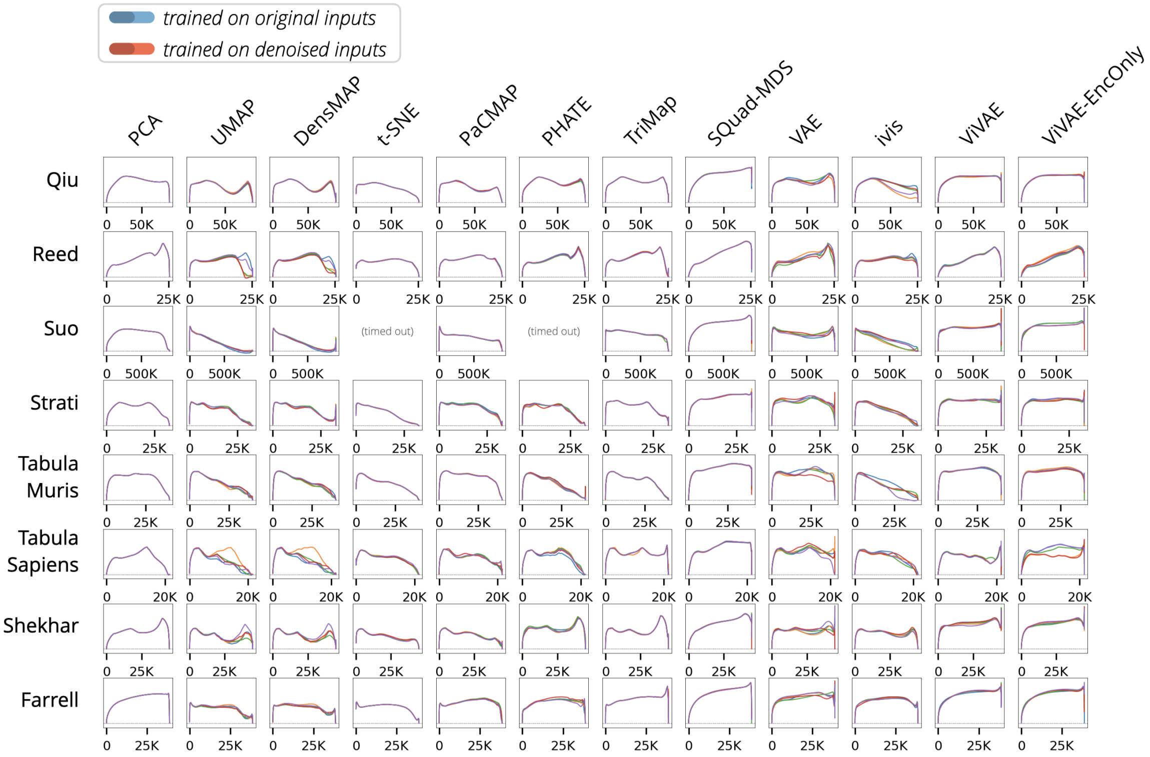


**Figure S2: Comparison of $R_{NX}$ curves for embeddings with and without the use of input denoising.** The $R_{NX}$ curves for each of 5 independent runs of each evaluated dimensionality reduction (DR) method (columns) on each of the evaluated datasets (rows) are shown. Results emanating from model training on non-denoised data are shown in different shades of blue, whereas those from model training on denoised data are shown in different shades of red. The *x*-axis shows neighbourhood scales. The *y*-axis shows $R_{NX}$ values.

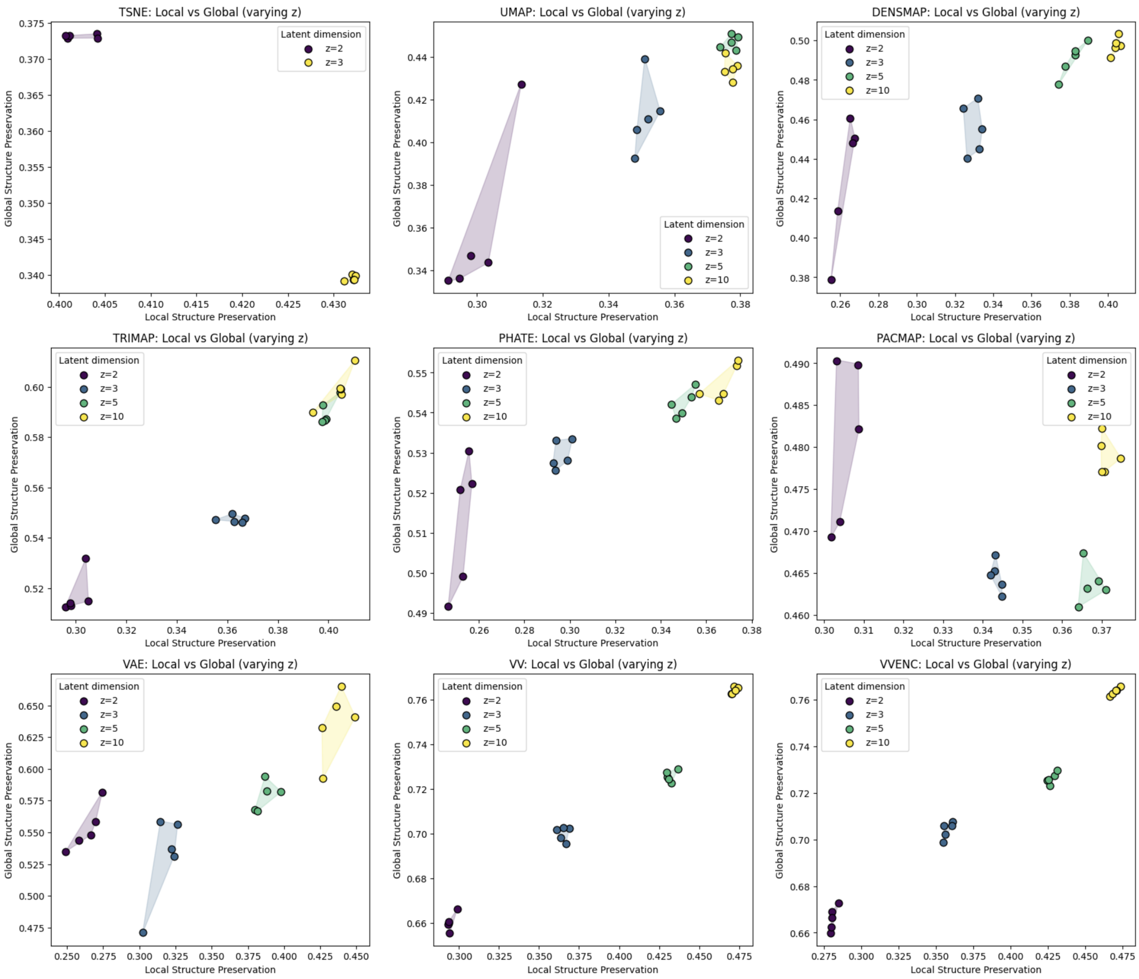


**Figure S3: Effects of varying embedding dimensionality on structure-preservation (SP) scores.** The Local SP ($x$-axis) and Global SP ($y$-axis) scores for embeddings of the *Farrell* dataset are shown per embedding method. Note that $t$-SNE only allows embedding dimensionality up to 3. Filled convex hulls are used to visualise the ranges of values between 5 independent runs with different random seeds per setting.

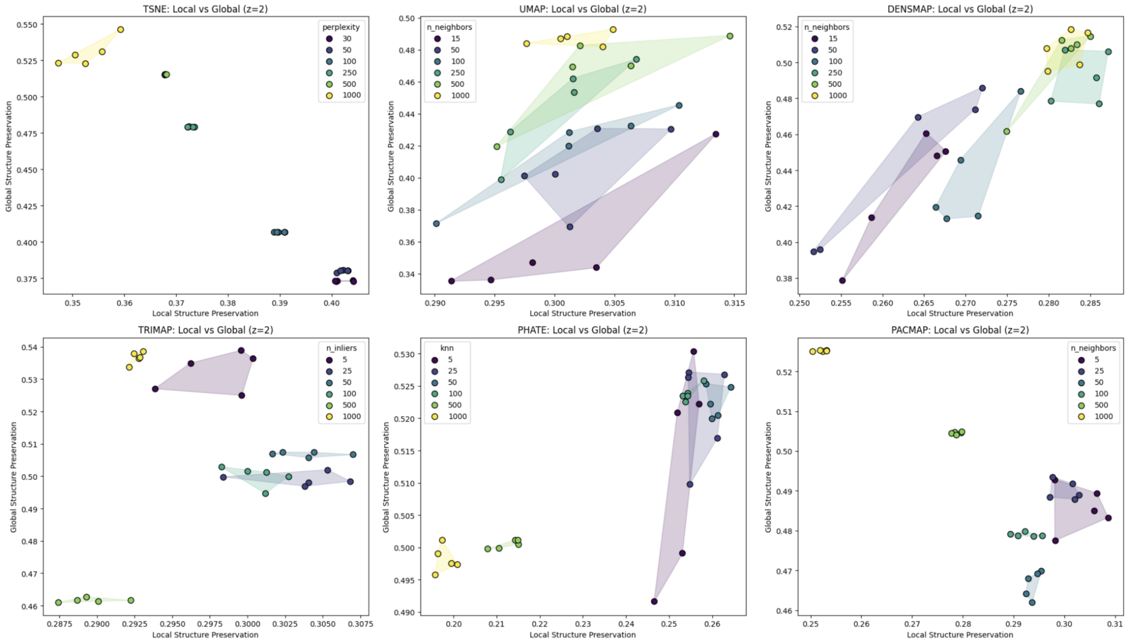


**Figure S4: Effects of varying the 'scale' hyperparameter on the structure-preservation (SP) scores.** The Local SP ($x$-axis) and Global SP ($y$-axis) scores for embeddings of the *Farrell* dataset are shown per embedding method. The name and range of tested values of the 'scale' hyperparameter per method are shown in respective subplot legends. Filled convex hulls are used to visualise the ranges of values between 5 independent runs with different random seeds per setting.

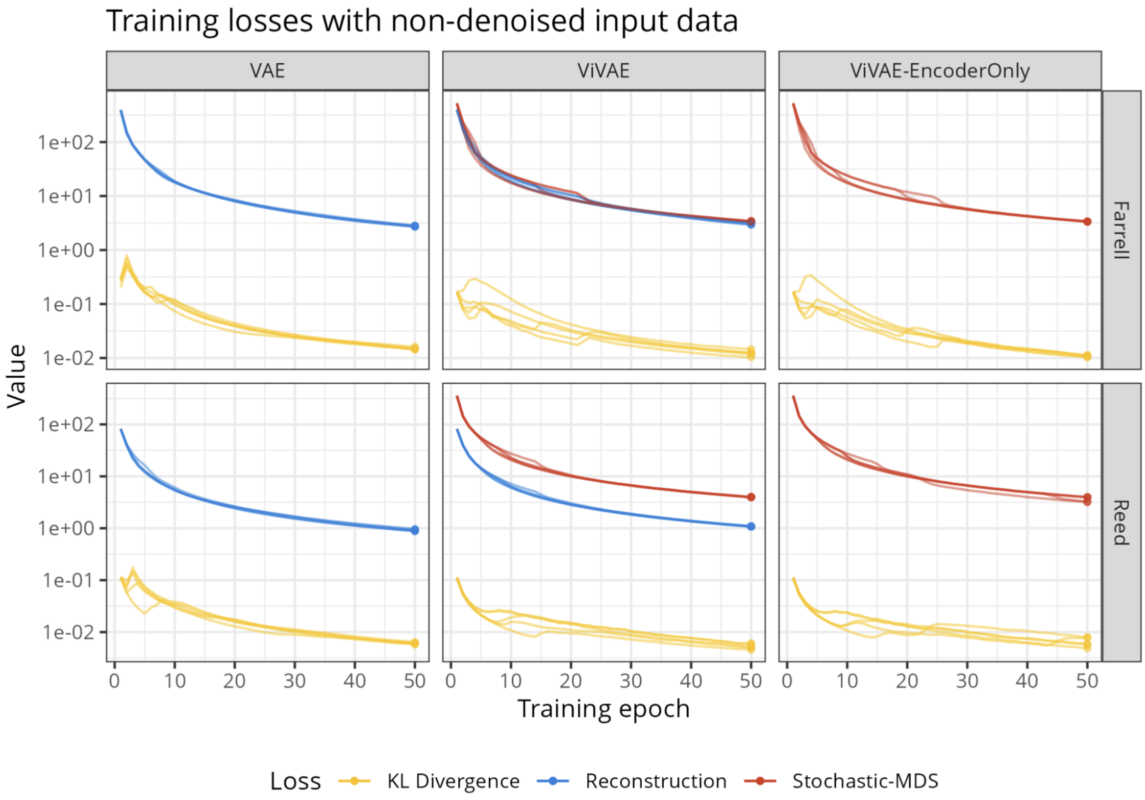


**Figure S5: Training loss across 50 training epochs for VAE, *ViVAE*, and *ViVAE-EncoderOnly*.** Values of reconstruction loss, KL-divergence, and stochastic-MDS loss per training epoch on non-denoised *Farrell* (developmental) and *Reed* (non-developmental) data are shown for a vanilla variational autoencoder (VAE; without stochastic-MDS) and the proposed models: *ViVAE* and *ViVAE-EncoderOnly*. Data from 5 independent runs are shown, using different shades of yellow, blue, and red for the respective loss terms. Defaultn and fixed architecture is used per model for comparability, with the *ViVAE-EncoderOnly* model lacking a decoder.

**Note S1**

**Algorithm 1** Denoising

**function** DENOISING($X, K, \lambda, n_i$)
▷ $X$: coordinate matrix, $K$: row-wise $k$-nearest-neighbour indices matrix, $\lambda$: shift coefficient, $n_i$: number of iterations
    $n \leftarrow \mathrm{rowCount}(X)$
    **for** $\mathrm{iter} \in 1..x_i$ **do**
        **for** $i \in 1..n$ **do**
            $\mathrm{idcs} \leftarrow K_{i*}$ ▷ neighbour indices
            $\mathrm{target} \leftarrow \mathrm{columnMeans}(X_{\mathrm{idcs},*})$ ▷ target coordinates
            $\mathrm{shift} \leftarrow \mathrm{target} - X_{\mathrm{i},*}$
            $X_{i*} \leftarrow X_{i*} + \lambda \mathrm{target}$
        **end for**
    **end for**
    **return** $X$
**end function**

**Note S2**

**Algorithm 2** Encoder Indicatrices

**function** CircleOnTangentPlane($x, h_1, h_2, r, n_p$)
▷ $x$: coordinates of point of origin, $h_1, h_2$: tangent vectors, $r$: indicatrix radius, $n_p$: number of polygon vertices to estimate indicatrix
  $h_1^n \leftarrow h_1 / L_2\text{norm}(h_1)$
  $h_2^n \leftarrow h_2 / L_2\text{norm}(h_2)$
  $s \leftarrow n_p^{-1}$ ▷ stepsize
  $P \leftarrow []$ ▷ points on circle
  **for** $i \in 0..(n_p - 1)$ **do**
    $a \leftarrow s \cdot 2\pi i$
    $P_i \leftarrow x + r\cos(a) h_1^n + r\sin(a) h_2^n$
  **end for**
  **return** $P$
**end function**

**function** EncoderIndicatrices($E, X, S, r, n_p$)
▷ $E$: trained encoder, $X$: row-wise input coordinates, $D$: indices of points for which to compute indicatrices
  $L_D \leftarrow E(X_D)$ ▷ 2-d embedding of selected points
  $Ind \leftarrow []$ ▷ list of embedded indicatrix points $\forall x \in X_D$
  **for** $i \in 1..\text{length}(D)$ **do**
    $d \leftarrow D_i$
    $J \leftarrow \text{jacobianMatrix}(\text{function} = E,\ \text{point} = L_d)$
    $U, S, V \leftarrow \text{singularValueDecomposition}(J)$ ▷ such that $J = U\text{diag}(S)V^T$
    $h_1 \leftarrow V_{*1}$ ▷ horizontal tangent vectors
    $h_2 \leftarrow V_{*2}$
    $P \leftarrow \text{CircleOnTangentPlane}(X_d, h_1, h_2, r, n_p)$
    $Ind_i \leftarrow E(P)$
  **end for**
  **return** $Ind$
**end function**

**Note S3**

**Algorithm 3** Extended Neighbourhood-Proportion-Error

**function** LIKENESSDISTRIBUTIONS($K, A$) ▷ $K$: row-wise $k$-nearest-neighbour indices matrix, $A$: vector of labels
  $n \leftarrow \mathrm{rowCount}(K)$
  $P \leftarrow \mathrm{unique}(A)$ ▷ populations
  $p \leftarrow \mathrm{length}(P)$
  $M \leftarrow [\,]_n$ ▷ point-level same-neighbour proportions
  $D \leftarrow [\,]_p$ ▷ population-level distributions of same-neighbour proportions
  **for** $i \in 1..n$ **do**
    $r \leftarrow K_{i*}$
    $M_i \leftarrow |A_r = A_i| / \mathrm{length}(A_r)$ ▷ point-level proportion of neighbours from same population
  **end for**
  **for** $j \in 1..p$ **do**
    $a \leftarrow |A = P_j|$
    $D_j \leftarrow \mathrm{histogram}(M_a)$ ▷ population-level distribution of same-neighbour counts
  **end for**
  **return** $D$
**end function**

**function** BASELINELIKENESSDISTRIBUTIONS($A$)
  $n \leftarrow \mathrm{rowCount}(K)$
  $P \leftarrow \mathrm{unique}(A)$
  $M \leftarrow [\,]_n$ ▷ same-neighbour proportions assuming random positions
  $p \leftarrow \mathrm{length}(P)$
  **for** $i \in 1..n$ **do**
    $M_i \leftarrow \mathrm{length}(A_i) / n$
  **end for**
  **for** $j \in 1..p$ **do**
    $a \leftarrow |A = P_j|$
    $D_j \leftarrow \mathrm{histogram}(M_a)$
  **end for**
**end function**

**function** xNPE($K_H, K_L, A$) ▷ $K_H$ and $K_L$: $k$-NN matrices for HD and LD data
  $P \leftarrow \mathrm{unique}(A)$
  $p \leftarrow \mathrm{length}(P)$
  $H \leftarrow$ LIKENESSDISTRIBUTIONS($K_H, A$)
  $L \leftarrow$ LIKENESSDISTRIBUTIONS($K_L, A$)
  $B \leftarrow$ BASELINELIKENESSDISTRIBUTIONS($A$)
  $U \leftarrow \mathrm{columnCount}(K_H)$ ▷ upper bound for EMDs
  $\delta \leftarrow [\,]_p$ ▷ population-level errors
  **for** $i \in 1..p$ **do**
    $\delta_i \leftarrow$ EARTHMOVER'SDISTANCE($H_i, L_i$)
    $\delta_i \leftarrow \delta_i / U$ ▷ rescaling
    $\delta_i \leftarrow \delta_i / B_i$ ▷ random baseline adjustment
  **end for**
  **return** $\delta$
**end function**

#### Note S4

```
import numpy as np
import scanpy as sc

hd = sc.read_h5ad('./scrnaseq.h5ad')

## If we filter by some condition (eg. tissue=='blood'):
# hd = hd[hd.obs['tissue']=='blood']
## If we are given raw counts:
# sc.pp.normalize_total(hd)
# sc.pp.log1p(hd)
sc.pp.scale(hd, max_value=10.)
sc.tl.pca(hd, svd_solver='arpack', n_comps=100)
pc = hd.obsm['X_pca']
np.save(f'pc.npy'), pc, allow_pickle=True)
```

### Note S5

#### Key resources table

| REAGENT or RESOURCE | SOURCE | IDENTIFIER |
|---|---|---|
| Deposited data | | |
| **Suo**: human multi-organ pre-natal T and NK cells scRNA-seq gene expression matrix | **CELLxGENE database**: https://cellxgene.cziscience.com/collections/b1a879f6-5638-48d3-8f64-f6592c1b1561 | Suo |
| **Strati**: human healthy bone marrow scRNA-seq gene expression matrix | **CELLxGENE database**: https://cellxgene.cziscience.com/collections/26b5b4f6-828c-4791-b4a3-abb19e3b1952 | Strati |
| **Reed**: human breast scRNA-seq gene expression matrix | **CELLxGENE database**: https://cellxgene.cziscience.com/collections/48259aa8-f168-4bf5-b797-af8e88da6637 | Reed |
| **Qiu**: murine embryonic T cells scRNA-seq gene expression matrix | **CELLxGENE database**: https://cellxgene.cziscience.com/collections/45d5d2c3-bc28-4814-aed6-0bb6f0e11c82 | Qiu |
| **TabulaMuris**: murine bone marrow scRNA-seq gene expression matrix | **CELLxGENE database**: https://cellxgene.cziscience.com/collections/0b9d8a04-bb9d-44da-aa27-705bb65b54ebBone marrow - A single-cell transcriptomic atlas characterizes ageing tissues in the mouse - 10x | TabulaMuris |
| **TabulaSapiens**: human thymus scRNA-seq gene expression matrix | **CELLxGENE database**: https://cellxgene.cziscience.com/collections/e5f58829-1a66-40b5-a624-9046778e74f5 | TabulaSapiens |
| **Farrell**: zebrafish embryos scRNA-seq gene expression matrix | **Single Cell Portal**: https://singlecell.broadinstitute.org/single_cell/study/SCP162/single-cell-reconstruction-of-developmental-trajectories-during-zebrafish-embryogenesis | Farrell |
| **Shekhar**: murine retina scRNA-seq gene expression matrix | **Single Cell Portal**: https://singlecell.broadinstitute.org/single_cell/study/SCP3/retinal-bipolar-neuron-drop-seq | Shekhar |
| Software and algorithms | | |
| ViVAE 1.0.0 | https://github.com/saeyslab/ViVAE | ViVAE |
| ViScore 1.0.0 | https://github.com/saeyslab/ViScore | ViScore |

| | | |
|---|---|---|
| scikit-learn 1.4.0 | https://pypi.org/project/scikit-learn/1.4.0/ | scikit-learn |
| scanpy 1.9.8 | https://pypi.org/project/scanpy/1.9.8/ | scanpy |
| umap-learn 0.5.5 | https://pypi.org/project/umap-learn/0.5.5/ | UMAP |
| SQuad-MDS | https://github.com/davnovak/SQuad-MDS.git | SQuad-MDS |
| TriMap 1.1.4 | https://pypi.org/project/trimap/1.1.4/ | TriMap |
| ivis 2.0.11 | https://pypi.org/project/ivis/2.0.11/ | ivis |
| PHATE 2.0.0 | https://pypi.org/project/phate/ | PHATE |
| PaCMAP 0.7.3 | https://pypi.org/project/pacmap/0.7.3/ | PaCMAP |
